# RNA recoding in cephalopods tailors microtubule motor protein function

**DOI:** 10.1101/2022.09.25.509396

**Authors:** Kavita J. Rangan, Samara L. Reck-Peterson

## Abstract

RNA editing is a widespread epigenetic process that can alter the amino acid sequence of proteins, termed ‘recoding’. In cephalopods, recoding occurs in most proteins and is hypothesized to be an adaptive strategy to generate phenotypic plasticity. However, how animals use RNA recoding dynamically is largely unexplored. Using microtubule motors as a model, we found that squid rapidly employ RNA recoding to enhance kinesin function in response to cold ocean temperature. We also identified tissue-specific recoded squid kinesin variants that displayed distinct motile properties. Finally, we showed that cephalopod recoding sites can guide the discovery of functional substitutions in non-cephalopod dynein and kinesin. Thus, RNA recoding is a dynamic mechanism that generates phenotypic plasticity in cephalopods and informs the functional characterization of conserved non-cephalopod proteins.

## Introduction

Adenosine to inosine (A-to-I) RNA editing occurs widely in animals and serves many functions, including the regulation and diversification of mRNAs (*1*). RNA editing in coding regions can generate nonsynonymous codon changes, termed ‘recoding’, resulting in protein variants with altered amino acid composition (*2, 3*). RNA recoding occurs extensively in soft-bodied cephalopods: ∼60% of all mRNAs are recoded in squid, octopus, and cuttlefish (*4, 5*). RNA sequencing has revealed tens of thousands of recoding sites across the transcriptomes of these animals, and most cephalopod transcripts harbor multiple recoding sites. Cephalopod RNA recoding therefore has the potential to dramatically diversify the proteome. It is hypothesized that cephalopods may use RNA recoding to dynamically modulate protein function in response to specific cellular (*6*) and environmental cues (*7, 8*). RNA editing varies across different cephalopod species and tissues (*4, 9*), but it is unknown if animals use differential recoding as a mechanism for phenotypic plasticity. For example, it is not known if RNA editing in cephalopods is responsive to environmental cues, or if protein variants generated through recoding in different tissues or environmental conditions display distinct functions. Thus, the endogenous roles of cephalopod RNA recoding remain largely unexplored.

Bioinformatic analyses of cephalopod RNA editing have led to a debate about the adaptive role and broad utility of recoding in these animals (*10*–*12*). The functional effects of recoding have only been assessed for a few cephalopod proteins, and most of this work has focused on voltage-gated potassium channels (*4, 7, 13*–*15*), which are also recoded in mammals (*16*). Outside of these proteins, the functional scope of cephalopod RNA recoding has not been explored. Furthermore, many cephalopod proteins are recoded at numerous sites, but it is unknown how multiple recoding sites within a single protein are employed in combination in different conditions to diversify protein function.

Many cephalopod recoding sites occur in uncharacterized amino acid residues that are widely conserved across other organisms, raising the possibility that cephalopod recoding sites could be harnessed as a unique map to reveal functional residues or targets of modulation in non-cephalopod proteins. This approach would be fundamentally different from conventional screening approaches, as recoding site substitutions represent natural variants that are likely non-deleterious and that animals may employ to enhance or modulate protein function in different contexts. To evaluate this idea, we first needed to determine if RNA recoding is indeed used to modulate protein function in cephalopods in different conditions, and also determine if individual cephalopod-guided recoding site substitutions in conserved residues alter protein function in non-cephalopod protein homologs.

Here, we set out to evaluate the function and utility of cephalopod recoding in the microtubule motor proteins dynein and kinesin. First, we wanted to determine if RNA recoding is used as a mechanism for phenotypic plasticity within animals. We identified differentially recoded variants of squid kinesin from different tissues and in response to changes in ocean temperature, a relevant environmental factor for these animals. We then investigated how recoded variants alter the motile properties of recombinant squid kinesin. Second, we wanted to determine if cephalopod recoding sites could reveal uncharacterized residues of functional significance in conserved non-cephalopod proteins. We assessed the functional effects of cephalopod recoding in conserved residues in yeast dynein and human kinesin.

## Results

### Microtubule motors proteins are extensively recoded in cephalopods

GO term analysis of edited and unedited proteins in the longfin inshore squid, *Doryteuthis pealeii*, revealed that proteins involved in transport and localization are significantly enriched amongst edited proteins (*4*) (fig. S1A, data S1). Across cephalopods, proteins involved in microtubule-based transport are extensively recoded 4 (fig. S1B and S1C, tables S1 and S2). Cytoplasmic dynein-1 (*DYNC1H1* encodes the dynein heavy chain in humans, “dynein” here) and kinesin-1 (*KIF5B* in humans, “kinesin” here) are ideal model proteins to characterize the effects of recoding on protein function. Both dynein and kinesin are highly conserved and are structurally and mechanistically well-characterized. In addition, single-molecule motility assays offer a robust method to quantify the effects of recoding on protein function (figs. S1D and S2).

### Tissue-specific recoding alters squid kinesin motility

If RNA recoding is a mechanism for phenotypic plasticity, then differential recoding would be expected to generate protein variants with distinct functions in different tissues or environmental conditions. We focused on RNA recoding of kinesin-1 from the squid genus *Doryteuthis*. We first asked how squid kinesin is differentially recoded in different tissues. RNA-sequencing indicates that the kinesin motor domain of *D. pealeii* contains 14 sites of RNA recoding (Fig. 1A) (*4, 5*), but it is unknown how these sites exist in combination in individual kinesin transcripts. We compared RNA recoding of the kinesin motor domain in two different squid neuronal cell populations: the stellate ganglion and the optic lobe (Fig. 1B). To identify what combinations of recoding sites are present in individual kinesin transcripts, we extracted RNA from the stellate ganglion and the optic lobe and then sequenced individual cDNA clones of the *D. pealeii* kinesin motor domain from each tissue. Transcripts from the stellate ganglion were more extensively recoded compared to the optic lobe (Fig. 1C), and we identified recoding site combinations that were unique to each tissue (Fig. 1D, fig. S3). In both tissues, we detected more multiply recoded transcripts than would be expected if each site were edited independently. We also detected significant tissue-specific pairwise correlations between recoding sites (Fig. 1E and F). These data indicate that RNA recoding occurs in unique combinations in different tissues to generate tissue-specific kinesin protein variants.

**Fig. 1.**
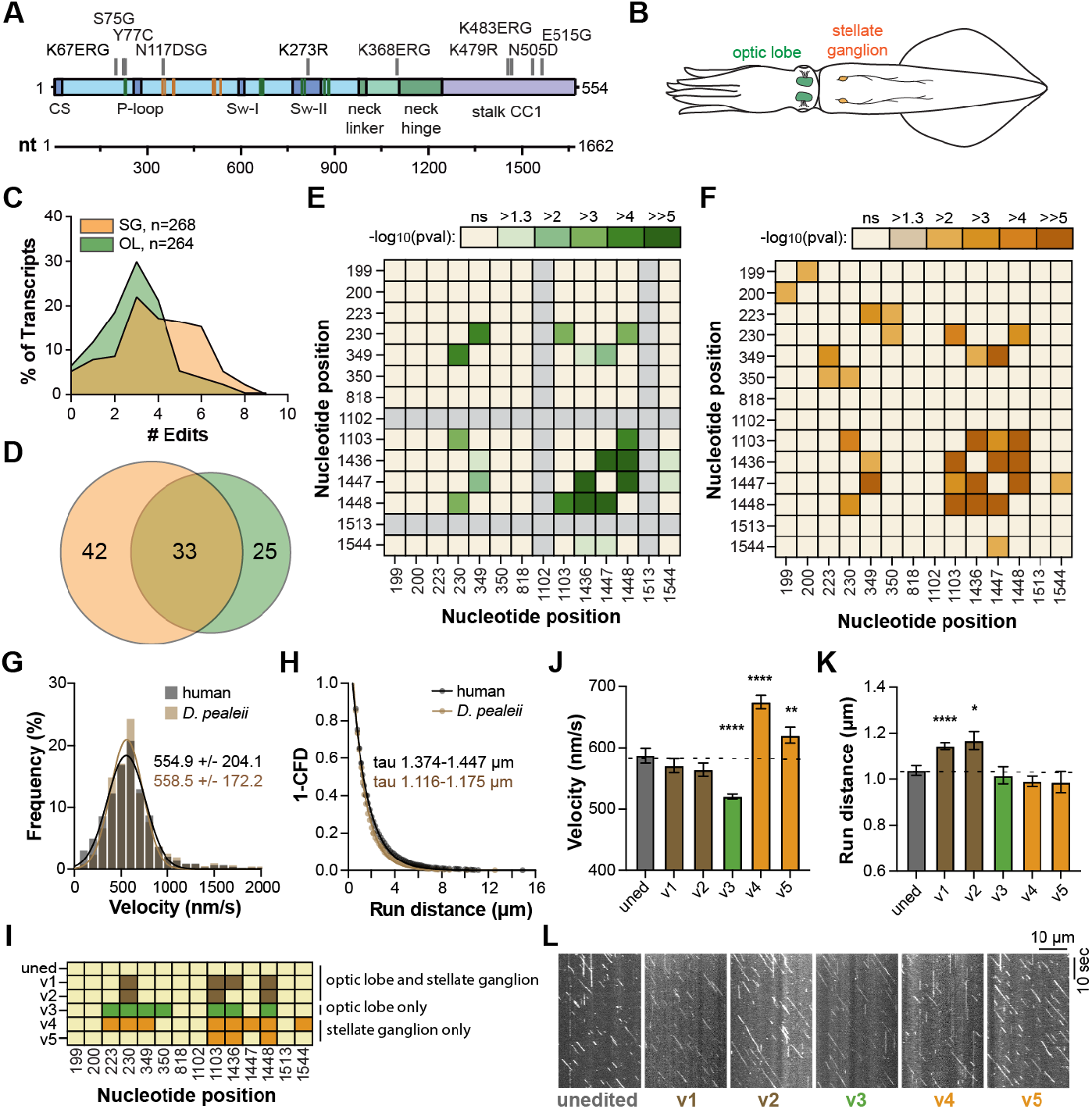
Tissue-specific recoding of squid kinesin generates functionally distinct motors. **(A)** Schematic indicating the location of recoding sites along the squid *D. pealeii* kinesin-1 motor domain (amino acids 1-554). The motor head is in blue, the neck is in green, and the stalk is in purple. A nucleotide (nt) ruler is included below. **(B)** Cartoon of *D. pealeii* with the optic lobes and stellate ganglion indicated. **(C, D)** Summary of clone sequencing of *D. pealeii* kinesin cDNA from the optic lobe (green) and stellate ganglion (orange). (C) Percent of transcripts detected with multiple edit sites and (D) number of unique transcripts detected in each tissue. See fig. S3 for details. **(E, F)** Pairwise correlations in recoding sites along the *D. pealeii* kinesin motor domain from the (E) optic lobe and (F) stellate ganglion. Gray squares indicate recoding sites that were not detected in any clones sequenced. Color indicates -log_10_ (p-value). **(G)** Velocity histograms (nm/s) of human kinesin (grey bars) and unedited *D. pealeii* kinesin (brown bars). The mean +/- SD of the Gaussian fit for each construct is indicated. (human kinesin (black line) R^2^ = 0.9773, n = 2432; *D. pealeii* kinesin (brown line) R^2^ = 0.9600 n = 1281). **(H)** Run distance analysis of human kinesin (grey circles) and unedited *D. pealeii* kinesin (brown circles). 1-cumulative frequency distributions were fit to a one-phase exponential decay. The 95% confidence interval of the mean decay constant for each construct is indicated as tau. (human kinesin (black line) R^2^ = 0.9971, n = 2191; *D. pealeii* kinesin (brown line) R^2^ = 0.9972 n = 1182). **(I)** *D. pealeii* recoding site variants generated in *D. pealeii* kinesin. Each variant contains a different set of A to G substitutions at the nucleotide positions indicated. Variants 1 and 2 were detected in both tissues (brown), variant 3 is specific to the optic lobe (green), and variants 4 and 5 are specific to the stellate ganglion (orange). **(J)** Mean velocities (nm/s) for each *D. pealeii* kinesin variant. **(K)** Mean decay constants (μm) of run distances for each *D. pealeii* kinesin variant. **(L)** Representative kymographs of *D. pealeii* kinesin variants in (J, K). See fig. S4 for frequency distributions of each *D. pealeii* kinesin variant. For all bar graphs of velocity measurements, velocity histograms were fit to a Gaussian, and bars represent the best-fit mean +/- 95% confidence intervals. For all bar graphs of run distance measurements, 1-cumulative frequency distributions were fit to a one-phase exponential decay to determine the mean decay constant, and bars represent the mean decay constant +/- 95% confidence intervals. Data were pooled from at least 2 independent protein preparations. For all bar graphs, p-values were calculated by two-tailed t-test, here comparing variants to unedited *D. pealeii* kinesin. Comparisons that are not significant are not labeled. See data S2 for velocity and run distance fit parameters including R^2^ and n, as well as exact p-values. Stars indicate the following p-values: ****, p < 0.0001; ***, p< 0.001; **, p<0.01; *, p<0.05; ns, p> 0.05

We next assessed the motile properties of different kinesin variants identified from the optic lobe and the stellate ganglion. The stellate ganglion includes the cluster of cell bodies that form the giant axon, which may necessitate distinct motor protein functions for rapid long-distance transport. We first generated an unedited GFP-tagged *D. pealeii* kinesin construct (*D. peal*K554), purified the recombinant protein, and quantified its motile properties in single-molecule motility assays. Unedited *D. pealeii* kinesin displayed similar velocities and modestly shorter run distances compared to human kinesin (*human*K560) (Fig. 1G and 1H). We generated five different recoded *D. pealeii* kinesin variants found in the optic lobe and stellate ganglion (Fig. 1I) and compared the motility of these variants to unedited *D. pealeii* kinesin. Recoded variants displayed significantly different motility characteristics compared to unedited *D. pealeii* kinesin (Fig. 1J-L, fig. S4, data S2). Variant 3, unique to the optic lobe, displayed decreased velocity, while variants 4 and 5, unique to the stellate ganglion, displayed increased velocity. Variants 1 and 2, which were detected in both tissues, displayed increased run distance, without altering velocity. Together, these data indicate that RNA recoding in different tissues generates unique kinesin variants with distinct motile properties. The increased velocities we observed in stellate ganglion-specific kinesin variants may reflect a need for faster long-distance axonal transport in these cells.

### Recoding of kinesin in squid is responsive to ocean temperature

Ocean temperature is a dynamic and ecologically relevant environmental factor for *Doryteuthis* squid. In the wild, these animals experience a broad range of temperatures during seasonal and daily depth migrations, frequenting waters as cold as 6 °C (*17, 18*). The motility of human kinesins changes dramatically with temperature *in vitro*; for example, human KIF5B (*19*) and KIF5A (*20, 21*) display decreased velocities and shorter run distances as temperature decreases. Unlike humans, cephalopods do not regulate their body temperature. We therefore wondered if temperature-dependent RNA recoding of kinesin could be one mechanism squid employ to modulate kinesin motility in different ocean temperatures. We hypothesized that RNA recoding could provide a homeostatic mechanism to maintain transport functions as these animals experience fluctuations in ocean temperature.

We examined the effects of ocean temperature on recoding of the kinesin motor domain of the opalescent inshore squid *Doryteuthis opalescens*. These animals make seasonal spawning migrations through the coast of San Diego, providing us local access to a squid population. The *D. opalescens* kinesin motor domain is 99.5% identical to *D. pealeii* and contains the same set of recoding sites. We first asked if RNA recoding of the *D. opalescens* kinesin motor domain changes in response to ocean temperature. *D. opalescens* egg casings were collected from the ocean, and hatchlings (Fig. 2A) were transferred to temperature-controlled seawater tanks for 24 hours before extracting RNA from individual whole animals (Fig. 2B). We exposed animals to water temperatures ranging from 6 °C to 20 °C, which encompasses the extreme range of temperatures these animals experience in the wild (*17, 18, 22*). We then sequenced kinesin from total cDNA to determine RNA editing levels at each individual recoding site along the kinesin motor domain from animals exposed to each temperature. For the majority of recoding sites in the kinesin motor domain, percent editing significantly increased as temperature decreased (Fig. 2C).

**Fig. 2.**
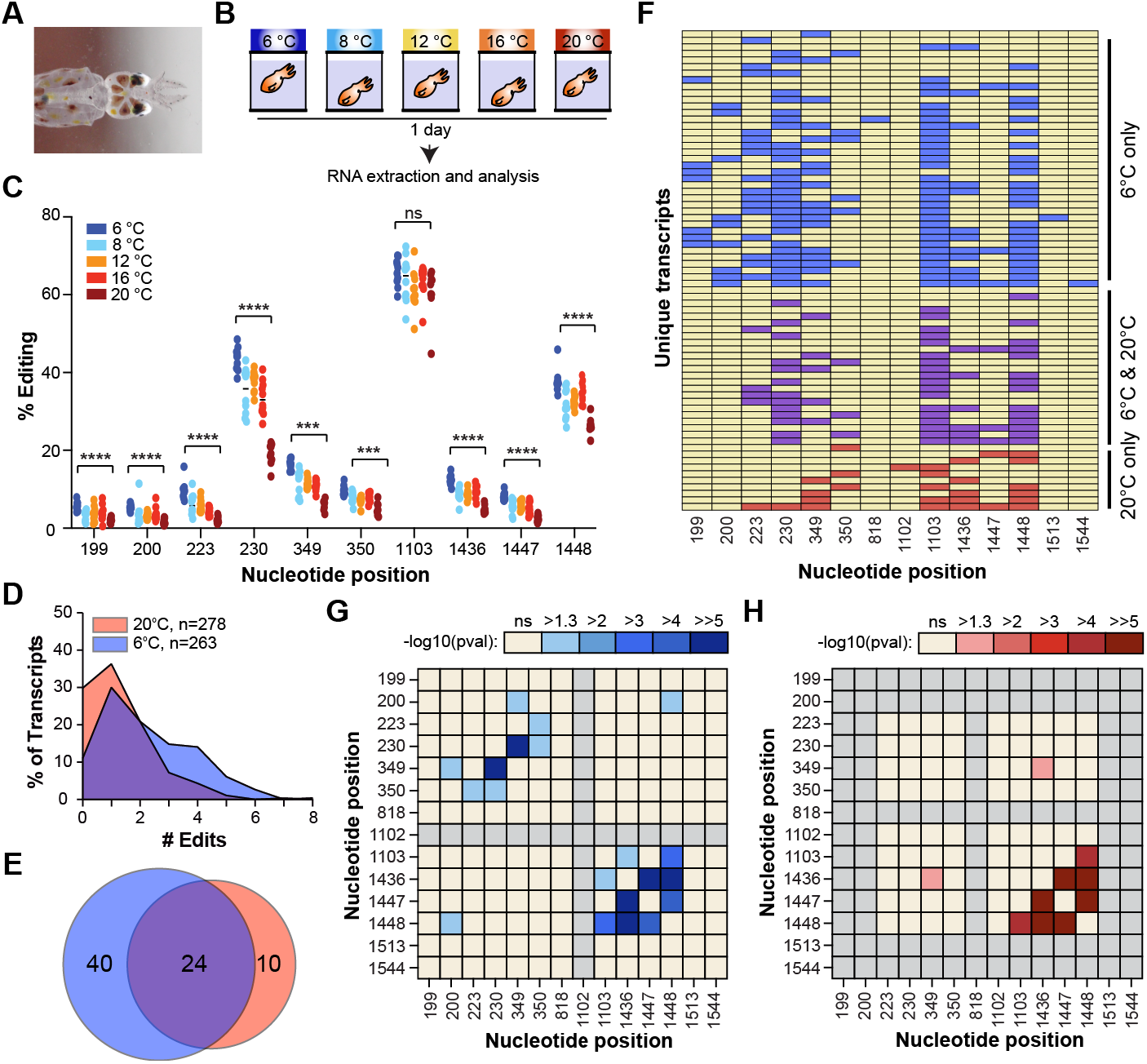
Squid kinesin transcripts are differentially recoded in response to ocean temperature. **(A)** Photograph of a *D. opalescens* hatchling. **(B)** Schematic of temperature assay. *D. opalescens* hatchlings were transferred to tanks of specified seawater temperatures for 24 hours, then collected for RNA extraction and analysis. **(C)** Percent editing at each recoding site along the *D. opalescens* kinesin-1 motor domain in individual animals exposed to different temperatures. Percent editing was measured by cDNA sequencing and peak height analysis at each site. Significance was determined by Mann-Whitney test. **(D, E)** Summary of clone sequencing of *D. opalescens* kinesin cDNA from animals exposed to either 6 °C (blue) or 20 °C (red) water. (D) Percent of transcripts detected with multiple edit sites and (E) number of unique transcripts detected at each temperature. See fig. S5 for details. **(F)** Unique recoding site combinations in *D. opalescens* kinesin detected from animals exposed to 6 °C (blue) or 20 °C (red) water. Each transcript (rows) has a different set of A to G substitutions at the nucleotide positions (columns) indicated. Combinations detected from both temperatures are in purple. **(G, H)** Pairwise correlations in recoding sites along the *D. opalescens* kinesin motor domain from animals exposed to (G) 6 °C or (H) 20 °C water. Gray squares indicate recoding sites that were not detected in any clones sequenced. Color indicates -log_10_ (p-value).

Next, we identified recoding site combinations in kinesin transcripts isolated from animals exposed to different seawater temperatures. We sequenced individual cDNA clones (representing individual kinesin transcripts) from animals exposed to either 6 °C or 20 °C seawater (Fig. 2D-F, fig. S5). Consistent with our bulk sequencing results, we observed higher editing levels in kinesin transcripts from animals exposed to 6 °C seawater as compared to 20 °C seawater (Fig. 2D). Sequencing individual transcripts allowed us to quantify the percentage of fully unedited transcripts; 30% of the transcripts sequenced from animals exposed to 20 °C seawater were unedited, as compared to 11% unedited transcripts in animals exposed to 6 °C seawater (Fig. 2D). In addition, transcripts from animals exposed to 6 °C were more extensively recoded, with 38% of transcripts harboring three or more recoding events as compared to 13% from animals exposed to 20 °C (Fig. 2D). Thus, RNA recoding of kinesin in squid is responsive to changes in ocean water temperature. Recoded variants of kinesin are preferentially generated in colder seawater, while unedited kinesin is preferred in warmer seawater. We also identified significant ocean temperature-specific pairwise correlations between recoding sites (Fig. 2G and 2H). This demonstrates that RNA recoding generates temperature-specific kinesin variants in response to environmental temperature. In addition, our data shows that temperature-dependent changes in RNA recoding occur rapidly in animals, within 24 hours.

### Recoding enhances squid kinesin motility in the cold

We hypothesized that *D. opalescens* kinesin variants that were more extensively recoded in cold seawater might display enhanced motility *in vitro* at cold temperatures compared to unedited *D. opalescens* kinesin. We generated an unedited *D. opalescens* kinesin construct (*Dopal*K554) and measured its motile properties in single-molecule motility assays at 8 °C and 25 °C (fig. S6). Similar to human kinesin (*21*), unedited *D. opalescens* kinesin exhibited slower velocities and shorter run distances in the cold (8 °C) as compared to 25 °C (Fig. 3A and 3B). We generated five different recoded *D. opalescens* kinesin variants, three of which were unique to animals exposed to cold seawater (Fig. 4C) and quantified the motile properties of these variants at 8 °C (Fig. 3D-G, fig. S7, data S2). Two cold-specific *D. opalescens* kinesin variants (variants 4 and 5) displayed significantly longer run distances at 8 °C compared to unedited *D. opalescens* kinesin, while slightly decreasing velocity. In addition, all the cold-specific *D. opalescens* kinesin variants displayed significantly increased microtubule landing rates (the frequency of kinesin binding to the microtubule) at 8 °C compared to unedited *D. opalescens* kinesin. Thus, acute changes in RNA recoding of squid kinesin in response to cold ocean temperature generate kinesin variants with enhanced motile properties in the cold. This may be a mechanism squid use to optimize kinesin activity and transport in different environments (Fig. 3H).

**Fig. 3.**
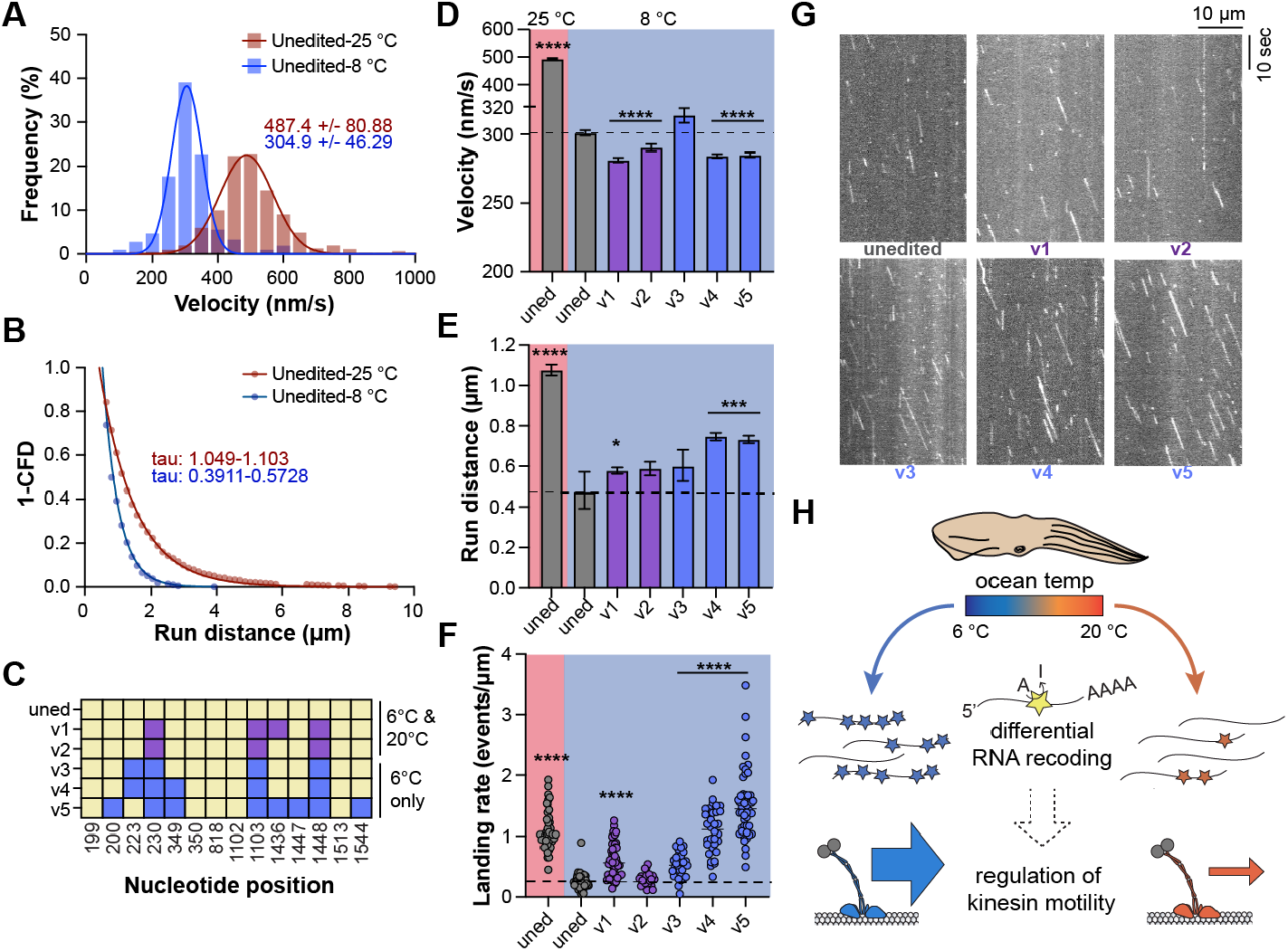
Cold-specific squid kinesin variants display enhanced motility in the cold. **(A)** Velocity histograms (nm/s) of unedited *D. opalescens* kinesin at 8 °C (blue bars) vs. 25 °C (red bars). The mean +/- SD of the Gaussian fit for each construct is indicated. (8 °C (blue line) R^2^ = 0.9873, n = 217; 25 °C (red line) R^2^ = 0.9794 n = 1186). **(B)** Run distance analysis of unedited *D. opalescens* kinesin at 8 °C (blue circles) and 25 °C (red circles). (8 °C (blue line) R^2^ = 0.9628, n = 217; 25 °C (red line) R^2^ = 0.9978 n = 1160). **(C)** *D. opalescens* recoding site variants generated in *D. opalescens* kinesin. Each variant contains a different set of A to G substitutions at the nucleotide positions indicated. Variants 1 and 2 were detected in animals exposed to both 6 °C and 20 °C (purple), and variants 3, 4, and 5 are specific to 6 °C (blue). **(D)** Mean velocities (nm/s) for each *D. opalescens* kinesin variant. Motility of each recoding variant at 8 °C was compared to unedited *D. opalescens* kinesin at 8 °C and 25 ° **(E)** Mean decay constants (μm) of run distances for each *D. opalescens* kinesin variant. **(F)** Landing rates (# of motors/μm of microtubule length) for each *D. opalescens* kinesin variant. For landing rate measurements, p-values were determined by two-tailed t-test with Welch’s correction. For (D-F), significance stars indicated are compared to unedited *D. opalescens* kinesin at 8 °C, and comparisons that are not significant are not labeled. **(G)** Representative kymographs of *D. opalescens* kinesin variants at 8 °C shown in (D-F). See fig. S7 for frequency distributions for each *D. opalescens* kinesin variant. **(H)** Model depicting how ocean temperature changes RNA recoding of squid kinesin-1 transcripts to specify kinesin motors with distinct motile properties.

**Fig. 4.**
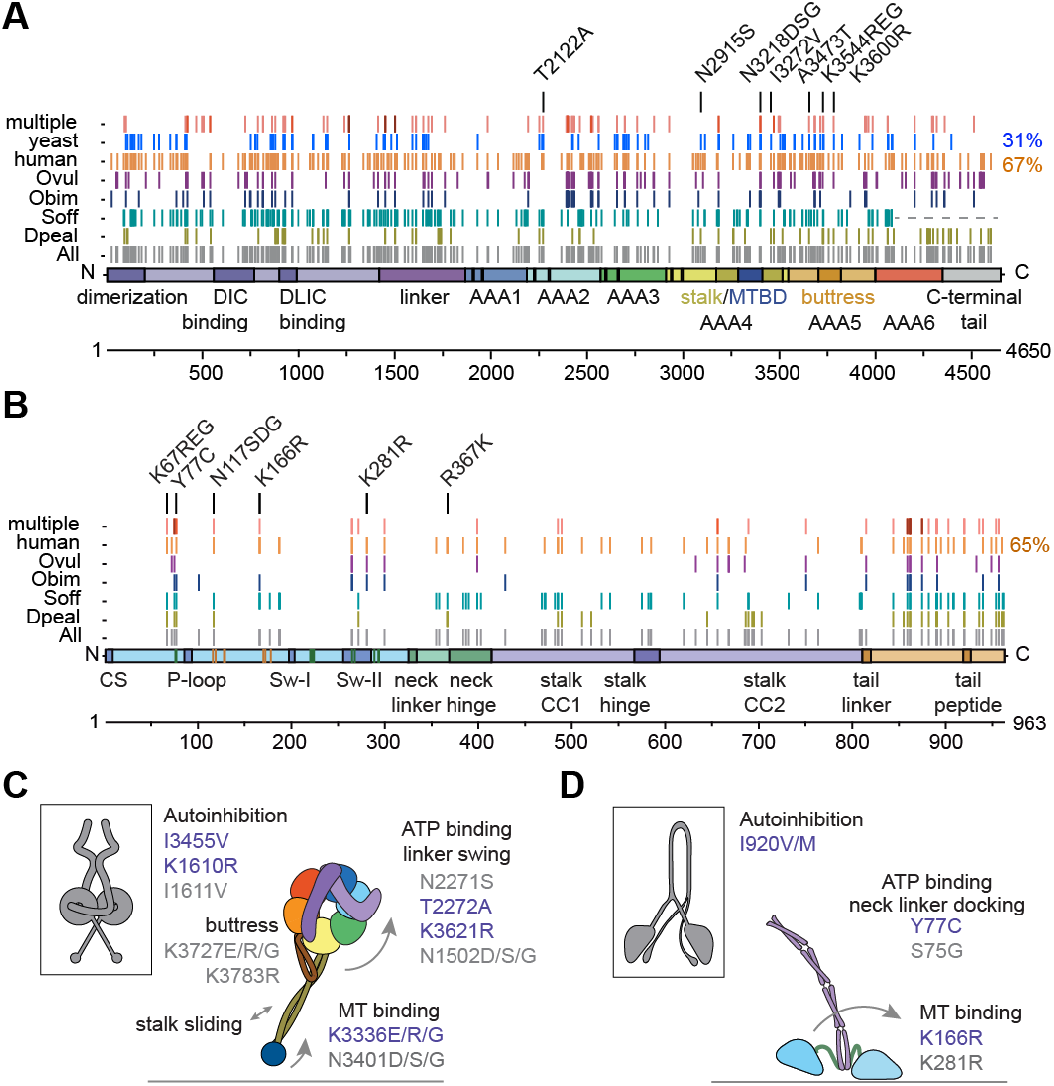
Cephalopod recoding sites occur in conserved residues in dynein and kinesin. **(A, B)** Cephalopod recoding sites distributed across (A) DYNC1H1 and (B) KIF5B. Domain features are indicated. (Multiple = sites detected in more than one cephalopod species, human = sites occurring in amino acids conserved with human DYNC1H1, yeast = sites occurring in amino acids conserved with *S. cerevisiae* DYN1, Ovul = *O. vulgaris* recoding sites, Obim = *O bimaculoides* recoding sites, Soff = *S. officinalis* recoding sites, Dpeal = *D. pealeii* recoding sites, All = combined cephalopod recoding sites.) Amino acid substitutions that we characterized in yeast GST-Dyn1(331kDa) dynein and human K560 kinesin are indicated. See data S3 and S4 for a list of recoding sites in dynein and kinesin respectively. The *S. officinalis Dync1h1* transcript assembly is truncated, as indicated by a dashed line **(C, D)** Schematics showing select cephalopod recoding sites associated with mechanochemical elements important for (C) dynein and (D) kinesin motility. Amino acids that have been previously implicated in each process are shown in blue, and sites that may be involved are shown in grey. Amino acid numbering is from human DYNC1H1 and human KIF5B.

### Recoding sites occur in conserved amino acids across dynein and kinesin

Our results with squid kinesin demonstrate that RNA recoding dynamically modulates protein function in different tissues and environmental conditions. We found that recoding generates a diversity of complex variants containing different combinations of recoded amino acids. Considered individually, many recoding sites occur in conserved amino acids; for example, in the squid kinesin motor domain, 7/10 recoded amino acids are conserved with human. We therefore sought to determine if individual recoding site substitutions in conserved amino acids could alter protein function in non-cephalopod proteins. We did so by characterizing the effects of cephalopod dynein and kinesin recoding site substitutions within well-studied homologs of these motors: yeast dynein and human kinesin.

We first surveyed individual cephalopod recoding sites across dynein and kinesin. We mined previously published transcriptome and editome data from four cephalopod species (*4*–*6*). Dynein and kinesin are highly conserved between these cephalopod species and human, sharing ∼74% and ∼65% amino acid sequence identity respectively. Both motors are heavily recoded in cephalopods, and recoding sites are widely distributed across conserved and non-conserved amino acids (Fig. 4A and 4B, data S3 and S4). Across the four cephalopod species we analyzed, there are 488 combined recoding sites in dynein, 82 of which are shared amongst multiple cephalopod species, and 143 combined recoding sites in kinesin, 37 of which are shared.

Dynein and kinesin are both dimeric motors that use the energy from ATP hydrolysis to generate conformational changes that ultimately drive unidirectional movement along the microtubule (*23*–*25*). Dynein’s motor domain contains a ring of AAA+ domains, and the nucleotide state at AAA1 and AAA3 govern the affinity between dynein’s microtubule binding domain (MTBD) and the microtubule. A long anti-parallel coiled-coil “stalk” and a shorter coiled-coil “buttress” connect dynein’s AAA+ ring to the MTBD, while the “linker” connects dynein’s AAA+ ring to the cargo binding end of the motor and makes large movements during dynein’s ATP hydrolysis cycle that generate force (*26*–*28*). Prior to activation, the dynein dimer exists in an autoinhibited “Phi” conformation that involves contacts between dynein’s stalk, AAA+ domains and linker (*29*). Kinesin’s motor domain is much smaller and more compact compared to dynein, with its single ATP binding and microtubule binding sites in close proximity (*30*). The motor domain of kinesin is connected to its coiled-coil dimerization and cargo binding domains via the “neck-linker”, which undergoes cycles of docking and undocking from the motor domain in a nucleotide-dependent manner (*31*). Kinesin also has an autoinhibited state, where its tail domain interacts with the motor (*32, 33*).

Some recoding sites occur in conserved amino acids within known mechanical elements of dynein and kinesin, but the specific substitutions generated through recoding at these sites have not been explored (Fig. 4C and D). For example in dynein, amino acid contact sites between the motor domain and the linker are recoded (*26*–*28*), as are contact sites important for stabilizing the auto-inhibited Phi form of dynein (*29*). In kinesin, amino acids involved in neck linker docking (*34, 35*), microtubule binding (*36*), as well as tail peptide autoinhibition (*32*) are also recoded. To our knowledge, only two recoding sites across the combined 513 sites in dynein and kinesin overlap with amino acids previously associated with disease (*37*), and recoding substitutions differ from disease-associated mutations at these sites. The vast majority of recoding sites in both dynein and kinesin occur in amino acids that have not been previously highlighted as functionally important. Thus, cephalopod recoding sites point to residues and substitutions that are not readily predicted by other methods.

By focusing on amino acid residues in yeast dynein and human kinesin that are conserved with and recoded in cephalopods, we dramatically reduce the number of substitutions to evaluate. For example, ∼1265 residues are conserved between human, yeast, and cephalopod dynein, but only 126 of these residues are also recoded. In addition, recoding guides what specific substitutions to evaluate. These substitutions represent natural variants found in cephalopods, and animals may employ either the recoded or non-recoded residue in different contexts. We chose to assess the effects of recoding site substitutions in well-characterized minimal motor constructs; for yeast dynein, we used a truncated, GST-dimerized construct (*38*) (GST-Dyn1(331kDa), “yeast dynein”) and for human kinesin, we used a truncated, GFP-tagged construct (*39*) (K560, “human kinesin”). We further narrowed a list of substitutions to evaluate by focusing on sites conserved within the motor domains of these minimal constructs that are recoded in at least 4% of RNAs from RNA-sequencing data (*4*–*6*) of any one cephalopod species. Applying these criteria, we generated a panel of 11 recoding site substitutions across seven amino acids in yeast dynein (Fig. 4A) and 10 recoding site substitutions across six amino acids in human kinesin (Fig. 4B).

For each substitution, we quantified velocity, run distance, and fraction of processive events in single-molecule motility assays. The majority of amino acid substitutions we assessed in both yeast dynein and human kinesin significantly altered motility (Fig. 5A-F, fig. S8 and S9, data S2). Unlike disease-associated mutations in dynein and kinesin (*40*–*42*), none of the cephalopod-guided substitutions we made abolished processivity, and most significantly increased run distance or velocity. Most of the substitutions we analyzed occurred in previously uncharacterized amino acids, indicating that cephalopod recoding sites can indeed be used as a guide to identify conserved residues of functional importance. We highlight examples in both motors below. For a discussion of all the substitutions we characterized, please refer to the supplementary text.

**Fig. 5.**
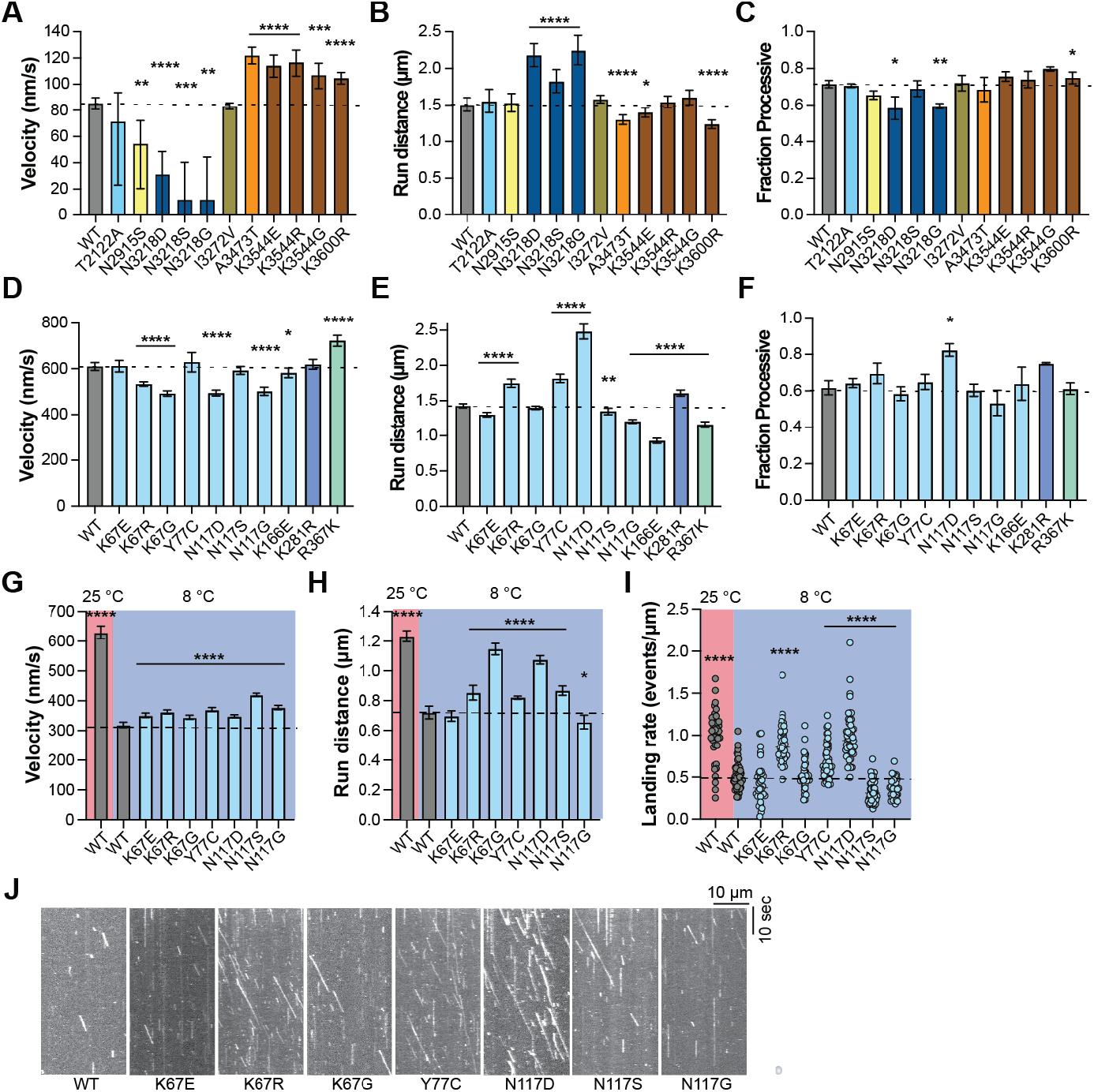
Recoding site substitutions reveal functionally important residues in yeast dynein and human kinesin. **(A)** Mean velocities (nm/s) for each yeast dynein mutant. **(B)** Mean decay constants (μm) of run distances for each yeast dynein mutant. **(C)** Fraction processive for each yeast dynein mutant. For fraction processive measurements, bars represent mean +/- SEM and p-values were determined by two-tailed t-test. For (A-C), bar coloring indicates domains depicted in Fig 4A. See fig. S8 for representative kymographs and frequency distributions for each yeast dynein mutant in (A-C). **(D)** Mean velocities (nm/s) for each human kinesin mutant. **(E)** Mean decay constants (μm) of run distances for each human kinesin mutant. **(F)** Fraction processive for each human kinesin mutant. For (D-F), bar coloring indicates domains depicted in Fig. 4B. See fig. S9 for representative kymographs and frequency distributions for each kinesin mutant in (D-F). **(G)** Mean velocities (nm/s) for each human kinesin mutant at 8 °C compared to wild type human kinesin at 25 °C and 8 °C. **(H)** Mean decay constants (μm) of run distances for each human kinesin mutant at 8 °C compared to wild type human kinesin at 25 °C and 8 °C. **(I)** Landing rates for each human kinesin variant at 8 °C compared to wild type human kinesin at 25 °C and 8 °C. **(J)** Representative kymographs of human kinesin mutants at 8 °C shown in (G-I). See fig. S10 for frequency distributions for each human kinesin mutant shown in (G-J).

### Recoding site substitutions alter yeast dynein motility

In dynein, several recoding site substitutions occur in the buttress. Deletion of the buttress/stalk interaction loop decouples ATPase activity from microtubule binding (*26*) and abolishes motility (*43*), but there is limited insight into key buttress residues outside of the buttress/stalk interaction loop. Eight amino acids within the buttress are sites of cephalopod recoding, and four of these are conserved with yeast dynein. We characterized recoding site substitutions in two of these amino acids located along the buttress coiled coil. Lys3544 can be recoded in squid and cuttlefish to Glu, Arg, or Gly. We found that substitution of Lys3544 to any of these diverse amino acids significantly increased velocity of yeast dynein compared to wild type without markedly altering run distance (Fig. 5A and B). Another lysine residue, Lys3600, is recoded in squid to Arg. Lys3600Arg resulted in increased velocity and decreased run distance compared to wild type (Fig. 5A and B). None of the substitutions we assessed are predicted to disrupt coiled coil structure, and the effects of these substitutions in yeast dynein highlight unexplored conserved residues in the buttress that can significantly influence motility.

We also characterized recoding site substitutions in the microtubule binding domain (MTBD). Eight amino acids in the MTBD are recoded in cephalopods, and four of these are conserved with yeast dynein. We assessed the effects of three recoding site substitutions in Asn3218, which is located in CC2 of the MTBD near the stalk (*44, 45*) and is not predicted to directly contact the microtubule. Asn3218 can be recoded in cuttlefish and octopus to Asp, Ser, or Gly. Asn3218Asp, Asn3218Ser, and Asn3218Gly in yeast dynein all resulted in decreased velocity and increased run distance compared to wild type (Fig. 5A and B).

### Recoding site substitutions alter human kinesin motility

Next, we analyzed the effects of cephalopod recoding site substitutions on the motile properties of human kinesin. Fourteen amino acids in the motor head are recoded in cephalopods, and 10 of these are conserved with human. We assessed recoding site substitutions in five of these amino acids. Two of the residues we looked at, Lys67 and Asn117, are uncharacterized surface-exposed residues for which there is little functional insight. Lys67 is in alpha-1b and can be edited to Glu, Arg, or Gly in squid and cuttlefish. We found that Lys67Glu resulted in a modest decrease in run distance, while Lys67Arg increased run distance and decreased velocity (Fig. 5D and E). These effects suggest that Lys67 may be involved in stabilizing long-range electrostatic interactions with the microtubule, similar to the computationally-predicted role for Lys68 (*46*). Asn117 is located in alpha-2b and can be edited to Asp, Ser, or Gly in squid and cuttlefish. We found that Asn117Asp decreased velocity and markedly increased run distance, while Asn117Gly decreased velocity and decreased run distance (Fig. 5D and E). Thus, many new functional residues in dynein and kinesin were identified using cephalopod RNA recoding sites as a guide.

### Cold-preferred recoding sites enhance human kinesin function in the cold

Three amino acids (Lys67, Tyr77, and Asn117) that were preferentially recoded in squid kinesin transcripts from animals exposed to the cold (Fig. 2F) are conserved with human kinesin. We hypothesized that individual substitutions in Lys67, Tyr77, and Asn117 in human kinesin might enhance motility of human kinesin in the cold. Indeed, Lys67Arg, Tyr77Cys, and Asn117Asp substitutions in human kinesin increased run distance at 25 °C compared to wild-type (Fig. 5E), which is consistent with the increase in run distance we observed with cold-preferred squid kinesin variants at 8 °C. We assessed the motility of recoding site substitutions in Lys67, Tyr77, and Asn117 in human kinesin at 8 °C (Fig. 5G-J, fig. S10, data S2) and found that Lys67Arg, Lys67Gly, Tyr77Cys, Asn117Asp, and Asn117Ser substitutions all significantly increased run distance of human kinesin at 8 °C compared to wild-type human kinesin (Fig. 5H). Lys67Arg, Tyr77Cys, and Asn117Asp substitutions also displayed increased landing rates at 8 °C (Fig. 5I). We also observed that Asn117Ser and Asn117Gly substitutions in human kinesin increased velocity and decreased landing rate at 8 °C compared to wild-type human kinesin. (Fig. 5G-I). These data highlight how condition-specific cephalopod recoding sites can also reveal functionally important amino acids in non-cephalopod homologs.

This survey of recoding site substitutions in yeast dynein and human kinesin provides an example for how cephalopod recoding sites can be used to identify conserved amino acids important for protein function, and further highlights the role of RNA recoding in diversifying protein function. Our data also suggests that recoding sites can reveal functional substitutions in protein regions where structural and comparative genetic approaches alone offer limited insight.

## Discussion

Our results indicate that cephalopods use RNA recoding as a mechanism for phenotypic plasticity. We show that squid dynamically tune kinesin function through RNA recoding to support physiological needs and acclimate to changing environmental conditions. These data provide evidence supporting a long-standing hypothesis that RNA recoding is a mechanism for temperature acclimation (*7, 8, 47*). Temperature-dependent recoding of kinesin or other proteins may be an important component of temperature acclimation in other animals as well, such as *Drosophila* and hibernating mammals, where global temperature-dependent changes in RNA editing have been described (*48*–*51*).

Our analysis of squid kinesin transcripts motivates understanding the specificity, mechanisms, and regulation of combinatorial RNA editing. Edit sites are typically determined from short-read RNA sequencing, and thus for most transcripts it is unclear how edit sites occur in combination along the same transcript. We identified tissue-specific and temperature-specific recoding site combinations encoding kinesin variants with distinct motile properties. Our data also indicate that recoding is correlated between sites in a condition-specific manner. Emerging long-read RNA sequencing methods (*52, 53*) should offer an opportunity to widely assess combinatorial RNA editing in diverse cell-types and conditions.

Our survey of recoding site substitutions in yeast dynein and human kinesin highlights how the cephalopod editome can be used to interrogate the function of conserved proteins. Multiple protein sequence alignments provide a comparative genetic approach to infer regions of conserved function. Cephalopod RNA editing offers an epigenetic lens to further pinpoint functionally important substitutions in conserved residues, complementing insights gained by structure and homology alone. This approach may be especially useful in protein regions where a priori identification of amino acid residues important for function is challenging and could be used to identify amino acid substitutions that enhance or modulate protein function under different contexts. Thus, we propose that cephalopod recoding sites represent a general resource and guide for characterizing other proteins. For example, numerous disease-associated proteins are highly recoded in cephalopods (eg. Tau, Amyloid beta, LRRK2) and characterizing recoding sites may offer a fresh perspective into the regulation and function of these proteins.

## Acknowledgements

We wish to thank Aaditya Rangan at the Courant Institute of Mathematical Sciences for pairwise correlation analysis of combinatorial RNA editing, Donovan Ventimiglia for help designing and coding the kymograph analysis software, Phil Zerofski at the Scripps Institute for Oceanography at UC San Diego for collecting *D. opalescens* egg cases, Lily McCormick for help setting up tanks, and Joshua Rosenthal at the Marine Biological Laboratory for gifting *D. pealeii* tissue samples. We thank Andres Leschziner, Donovan Ventimiglia, Steve Flavell, Andrew Gordus, Jenna Christensen, Eva Karasmanis, and Aga Kendrick for feedback on the manuscript. This paper was typeset with the bioRxiv word template by @Chrelli: www.github.com/chrelli/bioRxiv-word-template

## Author contributions

KJR and SRP conceived of the study. KJR performed the experiments. KJR and SRP wrote the manuscript.

## Materials and Methods

### Mapping of edit sites

Edit sites and editing levels for *D. pealeii, S. officinalis, O. vulgaris*, and *O. bimaculoides* were from previously reported RNA-seq data (*4*–*6*). Alignments were performed using T-coffee multiple protein sequence alignments (*54*) to determine amino acid conservation at sites of cephalopod recoding. Cephalopod *DYNC1H1* homologues were aligned to *H. sapiens* (DYHC1_HUMAN), *M. musculus* (DYHC1_MOUSE), *S. cerevisiae* (DYHC_YEAST), *A. nidulans* (DYHC_EMENI), and *D. melanogaster* (DYHC_DROME). Cephalopod KINH homologues were aligned to *H. sapiens* (KINH_HUMAN), *M. musculus* (KINH_MOUSE), and *D. melanogaster* (KINH_DROME).

### Gene ontology (GO) term enrichment analysis

GO term enrichment analysis was performed using the Blast2GO toolkit in the OmicsBox suite (Biobam; Valencia, Spain). The *D. pealeii* transcriptome and list of edited proteins were previously reported (*4, 5*). p-values were determined by Fisher’s exact t-test.

### Strain generation

#### Yeast strain generation

All dynein constructs were generated in *S. cerevisiae* strain W303a. GST-Dyn1(331kDa)(*38*) was modified using PCR-based methods and transformed using the lithium acetate method. The following point mutations were generated using QuikChange site-directed mutagenesis (Agilent) and strains were verified by DNA sequencing: T2122A, N2915S, N3218D, N3218S., N3218G, I3272V, A3473T, K3455E, K3544R, K3544G, K3600R.

#### E. coli strain generation

All kinesin constructs were expressed in *E. coli* BL21-Codon Plus(DE3) (Agilent). All mutations were generated using QuikChange site-directed and multi-site directed mutagenesis (Agilent) and all constructs were verified by DNA sequencing. Human K560-GFP was a gift from the Vale lab. The following point mutations were generated in human K560-GFP: K67E, K67R, K67G, Y77C, N117D, N117S, N117G, K166E, K281R, R367K. To generate unedited *D. pealeii* K554-GFP, the first 554 amino acids of *D. pealeii* KIF5B was determined from the *D. pealeii* transcriptome (*4, 5*), synthesized as a codon-optimized gene block (IDT), and cloned into pET17b. The following mutation variants were generated in *D. pealeii* unedited K554-GFP: v1: Y77C, K368R, K479R, K483R; v2: Y77C, K368R, K483R; v3: S75G, Y77C, N117G, K368R, K479R, K483R; v4: S75G, Y77C, N117D, K368R, K479R, K483G, E515G; v5: K368R, K479R, K483R. The *D. opalescens* KIF5B sequence was determined by PCR cloning KIF5B from *D. opalescens* cDNA using *D. pealeii* KIF5B-specific primers. The first 554 amino acids of KIF5B in *D. opalescens* contains three SNPs as compared to *D. pealeii* (E20Q, V352E, A405T) so these three mutations were introduced to generate unedited *D. opalescens* K554-GFP. The following mutation variants were generated in *D. opalescens* unedited K554-GFP: v1: Y77C, K368R, K479R, K483R; v2: Y77C, K368R, K483R; v3: S75G, Y77C, K368R, K483R; v4: S75G, Y77C, N117D, K368R, K483R; v5: K67R, Y77C, N117D, K368R, K479R, K483G, E515G.

### Protein purification

#### Dynein constructs

Cultures of *S. cerevisiae* for protein purification were grown, harvested, frozen and purified as described previously (*38*). Briefly, liquid nitrogen-frozen yeast cell pellets were lysed by grinding with a chilled coffee grinder and resuspending in dynein lysis buffer (DLB: final concentration 30 mM HEPES [pH 7.4], 50 mM potassium acetate, 2 mM magnesium acetate, 1 mM EGTA, 10% glycerol, 1 mM DTT) supplemented with 0.1 mM Mg-ATP, 0.5 mM Pefabloc, 0.05% Triton and cOmplete EDTA-free protease inhibitor cocktail tablet (Roche)). The lysate was clarified by centrifuging at 264,900 × g for 1 hour at 4 °C. The clarified supernatant was incubated with IgG sepharose beads (GE Healthcare Life Sciences) for 1.5 hours at 4 °C. The beads were transferred to a gravity flow column, washed with DLB buffer supplemented with 250 mM potassium chloride, 0.1 mM Mg-ATP, 0.5 mM Pefabloc and 0.1% Triton, and with TEV buffer (10 mM Tris–HCl [pH 8.0], 150 mM potassium chloride, 10% glycerol, 1 mM DTT, 0.1 mM Mg-ATP and 0.5 mM Pefabloc). Constructs were labeled with 5 μM Halo-TMR (Promega) in the column for 10 minutes at room temperature and unbound dyes were washed with TEV buffer at 4 °C. Dynein was cleaved from IgG beads via incubation with 0.15 mg/mL TEV protease for 1 hour at 16 °C. Cleaved proteins were filtered by centrifuging with Ultrafree-MC VV filter (EMD Millipore) in a tabletop centrifuge at 4 °C, flash frozen in liquid nitrogen and stored at −80 °C.

#### Kinesin constructs

Human K560-GFP and squid K554-GFP constructs were purified from BL21-CodonPlus(DE3)-RIPL (Agilent) *E. coli*. Cultures were grown to OD 0.6-0.8 and expression was induced with 0.75 mM IPTG for 16 hours at 18 °C. Cultures were pelleted, washed in PBS, and then liquid nitrogen-frozen. Frozen pellets were resuspended in lysis buffer (50 mM Tris [pH 7.5], 250 mM sodium chloride, 1 mM magnesium chloride, 20 mM imidazole [pH 7.4]) supplemented with 0.5 mM Mg-ATP, 10 mM beta-mercaptoethanol, 1 mM Pefabloc, cOmplete EDTA-free protease inhibitor cocktail tablet (Roche) and 50 mg/ml lysozyme) and kept on ice for 30 minutes. Cultures were lysed by sonicating at 50% power for 30 cycles with 5 second pulses and 20 seconds of rest on ice. Lysates were clarified by centrifuging at 92,600 x g for 30 minutes at 4 °C. The clarified supernatant was incubated with Ni-NTA agarose beads (Qiagen) for 1 hour at 4 °C. The beads were transferred to a gravity flow column, washed with lysis buffer supplemented with 0.5 mM Mg-ATP, 10 mM beta-mercaptoethanol, and 1 mM Pefabloc, then eluted with elution buffer (50 mM Tris [pH 7.5], 250 mM sodium chloride, 1 mM magnesium chloride, 250 mM imidazole [pH 7.4]) supplemented with 0.1 mM Mg-ATP, and 10 mM beta-mercaptoethanol. Eluants were then desalted using a PD-10 column (GE Healthcare) and buffer exchanged into BRB80 (80 mM K+PIPES [pH 6.8], 2 mM magnesium chloride, 1 mM EGTA) supplemented with 0.1 mM Mg-ATP and 0.1 mM DTT. Sucrose was added to the elution to a final concentration of 10%, and eluants were flash frozen in liquid nitrogen and stored at −80 °C.

### TIRF microscopy

Imaging was performed with an inverted microscope (Nikon, Ti-E Eclipse) equipped with a 100× 1.49 N.A. oil immersion objective (Nikon, Plano Apo). The xy position of the stage was controlled by a ProScan linear motor stage controller (Prior). The microscope was equipped with a MLC400B laser launch (Agilent), with 405 nm, 488 nm, 561 nm and 640 nm laser lines. The excitation and emission paths were filtered using appropriate single bandpass filter cubes (Chroma). The emitted signals were detected with an electron multiplying CCD camera (Andor Technology, iXon Ultra 888). Illumination and image acquisition was controlled by NIS Elements Advanced Research software (Nikon). Temperature-controlled motility assays were performed using a Linkam PE100-NIF inverted Peltier stage and T96 System Controller. Flow chamber slides were covered with a custom copper-plated aluminum fitting and empirical temperature of the flow chamber was monitored using a Type K thermocouple sensor probe.

### Motility assays

Single-molecule motility assays were performed in flow chambers made with double-sided tape using the TIRF microscopy set-up described above. No. 1–1/2 coverslips (Corning) were used for the flow chamber assembly and sonicated in 100% ethanol for 10 min to reduce non-specific protein binding. Taxol-stabilized microtubules with ∼10% biotin-tubulin and either ∼10% Alexa488-tubulin or Alexa405-tubulin were attached to the flow chamber via biotin-BSA and streptavidin as described previously (*55*). For each frame, kinesin-GFP and TMR-dynein were exposed for 100 ms with the 488 nm laser and 561 nm laser, respectively.

For motility assays, a final concentration of 2.5-5 pM of dynein or 50-500 pM of kinesin was flowed into the flow chamber pre-assembled with taxol-stabilized microtubules. Motor concentrations used were empirically determined to optimize the number of runs per kymograph. The final imaging buffer contained DLB (30 mM HEPES [pH 7.4], 2 mM magnesium acetate, 1 mM EGTA, 10% glycerol, 50 mM potassium acetate), 20 μM taxol, 1 mM Mg-ATP, 1 mg/mL casein, 1 mM DTT, 71.5 mM β-mercaptoethanol and an oxygen scavenger system (0.4% glucose, 45 μg/ml glucose catalase, and 1.15 mg/ml glucose oxidase). Microtubules were imaged first by taking a single-frame snapshot. Dynein was imaged every 500 ms for 5 minutes, and kinesin was imaged every 200 ms for 1 minute. At the end, microtubules were imaged again by taking a snapshot to check for stage drift and images showing drift were omitted from analysis. Dynein was imaged for no longer than 20 minutes, and kinesin was imaged for no longer than 10 minutes. For motility assays performed at 25 °C and 8 °C, slides were equilibrated on the Peltier stage for 1.5 minutes before imaging and temperature was monitored immediately before and after imaging using a thermocouple probe.

### Kymograph analysis

Motility movies were analyzed using an analysis pipeline outlined in fig. S2. First, kymographs were generated using an ImageJ macro as previously described(*56*). Kymographs were then pixel classified using Ilastik 1.3.3post3(*57*), and pixel-classified images were analyzed with custom Matlab scripts (Matlab R2021a Update 4). Individual runs were parsed from the skeletonized images and velocity and run distance measurements were determined. Runs shorter than 4 frames were excluded from analysis. Processive runs were defined as any run with a velocity of 15 nm/s or greater. Because motor velocity can change throughout a run, velocity measurements reflect segment velocities. Segments were defined using the findchangepts function to detect changes in run mean and slope. Landing rates for each kymograph were calculated as total number of events (motile + non-motile)/μm of microtubule length.

#### Statistical analysis

All statistical analyses were determined using Prism9 (GraphPad). See data S2 for fits and p-values. For velocity measurements, mean velocities were determined by fitting velocity histograms to a Gaussian distribution and p-values comparing the best-fit values were calculated by two-tailed t-test. For run distance measurements, mean decay constants were determined by fitting 1-cumulative frequency distributions to a one-phase exponential decay and p-values comparing the best-fit values were calculated by two-tailed t-test. For percent processivity measurements, p-values were determined by two-tailed t-test. For landing rate measurements, p-values were determined by two-tailed t-test with Welch’s correction.

#### Code availability

Custom Matlab scripts used in kymograph analysis are available on GitHub (repository kjran/Kymo_analysis).

### Determination of edit site combinations from *D. pealeii* tissues

Stellate ganglion and optic lobes were a gift from the Rosenthal lab at the Marine Biological Laboratory. Samples were lysed and total RNA was extracted using the RNAeasy Plus extraction kit (Qiagen) as per the manufacturer’s protocol. cDNA was generated using Accuscript High-fidelity 1st-strand cDNA synthesis kit (Agilent). The kinesin-1 motor domain was PCR amplified from cDNA using gene-specific primers and cloned into the pCR-XL-2-TOPO vector (Invitrogen) as per the manufacturer’s protocol. Individual clones were selected and sequenced to determine edit site combinations of individual transcripts. For each data set, three samples were processed and 100 clones for each sample were sequenced. Sequences with incomplete or low-quality sequencing for any portion of the motor domain were omitted and data were pooled per condition. Correlation analysis of recoding sites was performed as follows: The observed instances of each recoding site pair were compared against all possible outcomes for each pair given the number of transcripts sequenced. The p-value for each pairwise comparison was determined by calculating the probability of observing a more rare scenario given the null hypothesis that transcript sequences are independent of one another and recoding at each site along a transcript occurs independently at the observed frequency.

### *D. opalescens* temperature experiment

*Doryteuthis opalescens* egg cases were collected off the Scripps Pier in San Diego, CA. Water temperature at depth was 12-13 °C. Egg casings were transferred to seawater tanks maintained at 12-13 °C. Tanks were monitored for temperature stability. Newly hatched animals (less than 24 hours old) were transferred to tanks maintained at 6 °C, 8 °C, 12 °C, 16 °C, or 20 °C and held at these temperatures for 24 hours. Animals were then flash frozen in liquid nitrogen and stored at −80 °C.

### Determination of editing levels and edit site combinations from D. *opalescens*

RNA extraction, cDNA synthesis, and PCR amplification of kinesin was performed as described above for 10 animals from each temperature. PCR products were directly sequenced using reverse primers, and percent editing at each recoding site was determined by peak height analysis calculated by (C peak height)/(C + T peak heights). Only high quality, complete sequencing data was used for percent editing analysis. Significance between 6 °C and 20 °C editing levels was determined by non-parametric Mann-Whitney test and the false discovery rate was determined by applying the corrected method of Benjamini and Yekutieli. Edit site combinations were determined as described above for three animals exposed to 6 °C water, and three animals exposed to 20 °C water.

## Supplementary Text

### Description of recoding site substitutions in yeast dynein not described in the main text (refer to Fig. 5A-C)

#### Thr2122

Thr2122 is located within the H2 insert loop in AAA2L that contacts the linker when the linker bends upon ATP binding at AAA1 and ring closure (*28*). A triple mutant of this contact region containing T2122G in yeast GST-Dyn1(331kDa) displays decreased velocity in single molecule motility assays (*58*) and deletion of the entire H2 insert region in *Dictyostelium* dynein results in lower microtubule-stimulated ATPase activity (*26*). In our assays, Thr2122Ala did not significantly affect velocity or run distance. However, multiple amino acids in the H2 hairpin insert are recoded in cuttlefish (hTyr2265 (recoded to Cys), hAsn2271 (recoded to Ser), hThr2272 (recoded to Ala)) and these in other assays or in combination may affect linker docking at AAA2.

#### Asn2915

This site is edited in *D. pealeii*. Asn2915 is located at the edge of AAA4L near the AAA3/4 interface. Asn2915Ser modestly decreased velocity without altering run distance.

#### Ile3272

This site is edited in *D. pealeii*. Ile3272 is located in the stalk and is an important stalk/stalk contact site for stabilizing the autoinhibited Phi form of dynein (*29*). An Ile3272Ala mutation in full-length yeast dynein results in increased run distance and decreased velocity (*59*). In our assays, Ile3272Val did not significantly affect velocity or run distance.

#### Ala3473

Both cephalopods and human encode a threonine at this position while *S. cerevisiae* encodes an alanine. *D. pealeii* and cuttlefish edit this position to an alanine, and editing occurs to almost 100% in the *D. pealeii* giant axon. We found that Ala3473Thr increased velocity and decreased run distance. Ala3473 lies at the edge of an exposed loop within AAA5L that is near contact sites between AAA5 and the linker (*27*), and a known contact site between AAA5 and the linker is also a cuttlefish recoding site (hLys3621 (recoded to Arg)) as is the adjacent residue (hAsn3622 (recoded to Asp)). Mutation of linker contact sites in AAA5 decreases single molecule velocity of GST-Dyn1(331kDa) (*27*). Another residue (yArg3476) lies on the opposite side of the AAA5 loop containing Ala3473 and may be involved in stabilizing electrostatic interactions between AAA5/AAA5 domains in the Phi conformation; a Arg3476Asp mutation in full-length yeast dynein results in increased run distance (*59*).

### Description of recoding site substitutions in human kinesin not described in the main text (refer to Fig. 5D-F)

#### Tyr77

This site is predicted to help stabilize formation of the cover neck bundle (CNB) and neck linker docking (*34*), and an aromatic residue at this position is highly conserved across kinesins (*35*). The analogous site in Kif1A (Tyr89) is mutated to aspartic acid in hereditary spastic paraplegia (HSP), and in single molecule motility assays Tyr89Asp results in slower velocity and increased run distance (*35*). Tyr77 is edited to a cysteine in cuttlefish (*S. officinalis*), octopus (*O. bimaculoides*), and squid (*D. pealeii*), and a Tyr77Cys mutation in human K560 resulted in increased run distance without altering velocity. Other mutations that destabilize CNB formation and neck linker docking have been shown to increase run distance in single molecule motility assays while decreasing force generation and motility of high-load cargo in cells (*60*). Tyr77 is almost 100% edited to cysteine in the squid giant axon, which may reflect a need for long distance transport in this compartment.

#### Lys166 and Lys281

These sites are located along the microtubule interaction face of the motor domain, and mutation of these residues to alanine results in decreased microtubule binding (*36*). Lys281Ala has a more severe effect on microtubule binding compared to Lys166Ala, and mutation of an adjacent residue (Arg280) to serine in HSP also results in decreased microtubule binding affinity (*61*). In *O. bimaculoides* and *O. vulgaris*, Lys281 is recoded to arginine, which would be expected to strengthen the electrostatic interaction between the motor head and the microtubule. In contrast, in cuttlefish, Lys166 can be recoded to glutamic acid, which would be expected to weaken the interaction with the microtubule. (Lys166 can also be recoded to an Arg and Gly but we did not characterize these substitutions.) We found that Lys281Arg increased run distance while Lys166Glu decreased run distance, which is consistent with the proposed role of these residues in stabilizing electrostatic interactions with the microtubule. Neither of these substitutions markedly affected velocity.

#### Arg367

This site is in the neck coiled coil. Humans encode an arginine at this position, while cephalopods encode a lysine. In squid, Lys367 is almost 100% edited to Arg. An Arg367Lys mutant in K560-GFP resulted in increased velocity and decreased run distance.

---

**Fig. S1.**
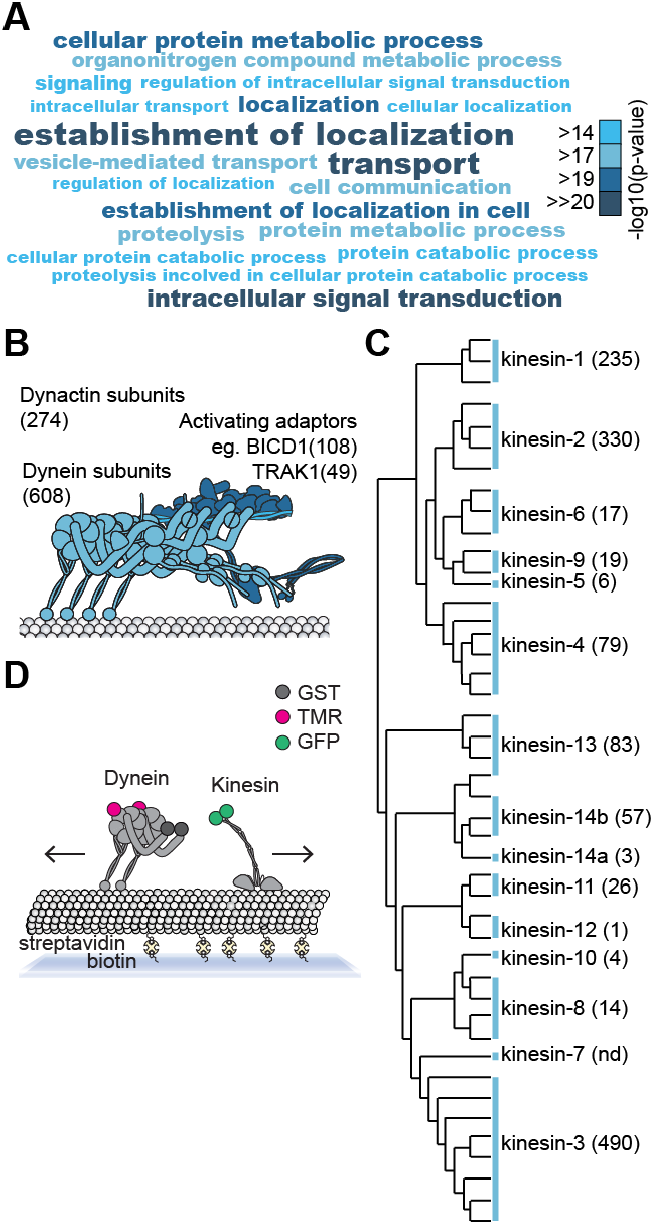
Microtubule motor proteins present an ideal model for exploring the function of cephalopod recoding. **(A)** Top 20 GO terms (biological process) significantly enriched amongst *D. pealeii* edited transcripts as compared to the full *D. pealeii* transcriptome. Shade indicates -log_10_ (p-value). See data S1 for details. **(B, C)** Sum of recoding sites from *D. pealeii, S. officinalis, O. bimaculoides*, and *O. vulgaris* across (B) dynein transport complex proteins and (C) kinesin family proteins. Kinesin phylogenetic tree reproduced from (Hirokawa et al., 2009). See tables S1 and S2 for details. (**D)** Schematic of a single-molecule motility assay. Biotinylated, fluorescently labeled microtubules are attached to a coverslip and motility of fluorescently labeled motor proteins is measured by total internal reflection fluorescence (TIRF) microscopy.

**Fig. S2.**
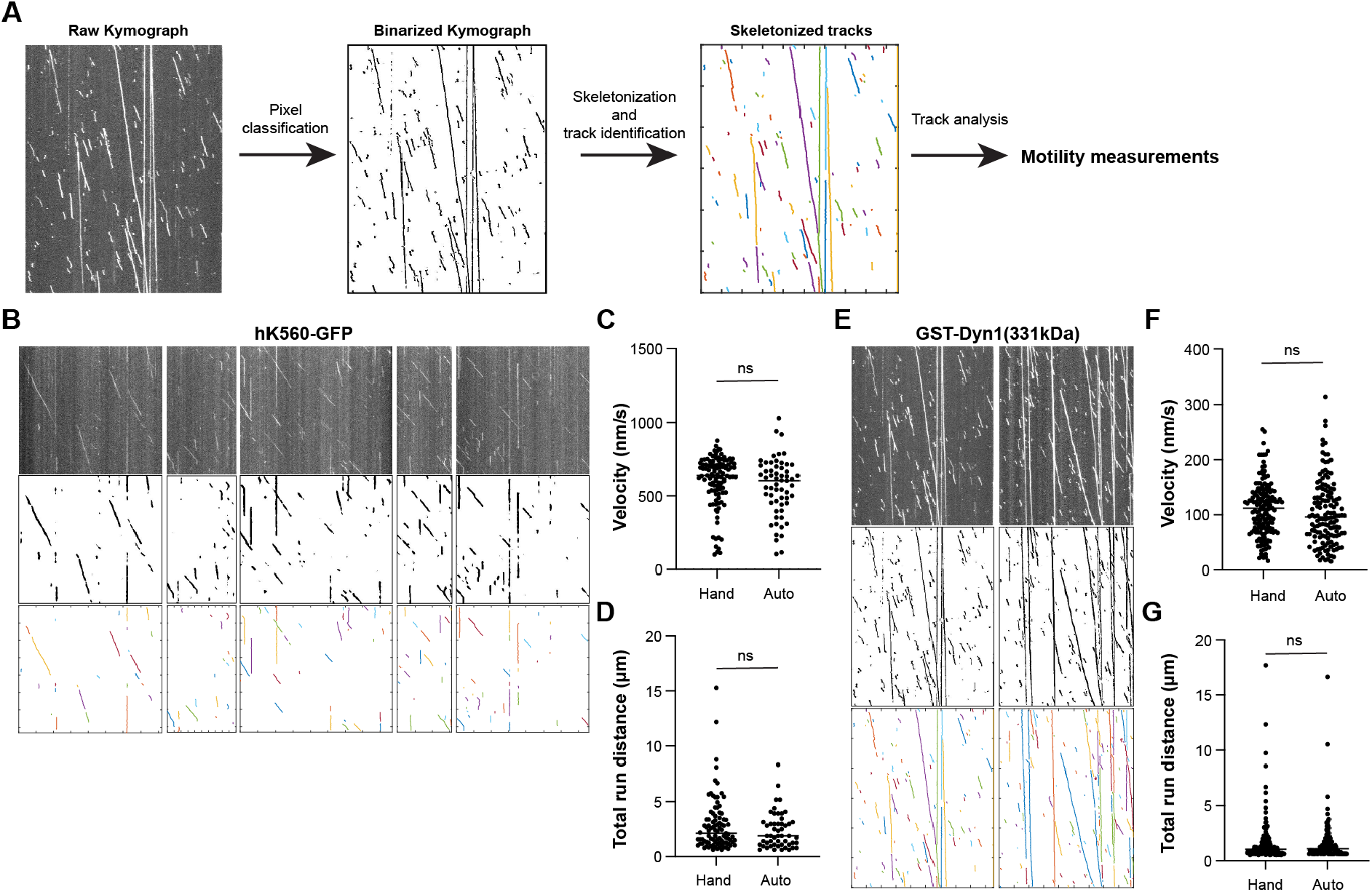
Comparison of custom automated kymograph analysis with hand-tracked kymograph analysis. **(A)** Automated kymograph analysis pipeline. See Methods and Github for details. Kymographs were binarized using Ilastik pixel classification. Binarized kymographs were skeletonized, and individual tracks were identified and analyzed for motility parameters including velocity and run distance using custom Matlab scripts. **(B)** Kymographs of human kinesin (K560) alongside the pixel classified and skeletonized kymograph images. These kymographs were used for directly comparing hand-tracking vs. automated tracking in (C-D). **(C)** Individual velocity measurements (nm/s) of human kinesin as determined by hand-tracking vs. automated tracking. **(D)** Individual run distance measurements (μm) of human kinesin as determined by hand-tracking vs. automated tracking. **(E)** Kymographs of yeast dynein (GST-Dyn1(331kDa)) alongside the pixel classified and skeletonized kymograph images. These kymographs were used for directly comparing hand-tracking vs. automated tracking in (F-G) **(F)** Individual velocity measurements (nm/s) of yeast dynein as determined by hand-tracking vs. automated tracking. **(G)** Individual run distance measurements (μm) of yeast dynein as determined by hand-tracking vs. automated tracking.

**Fig. S3.**
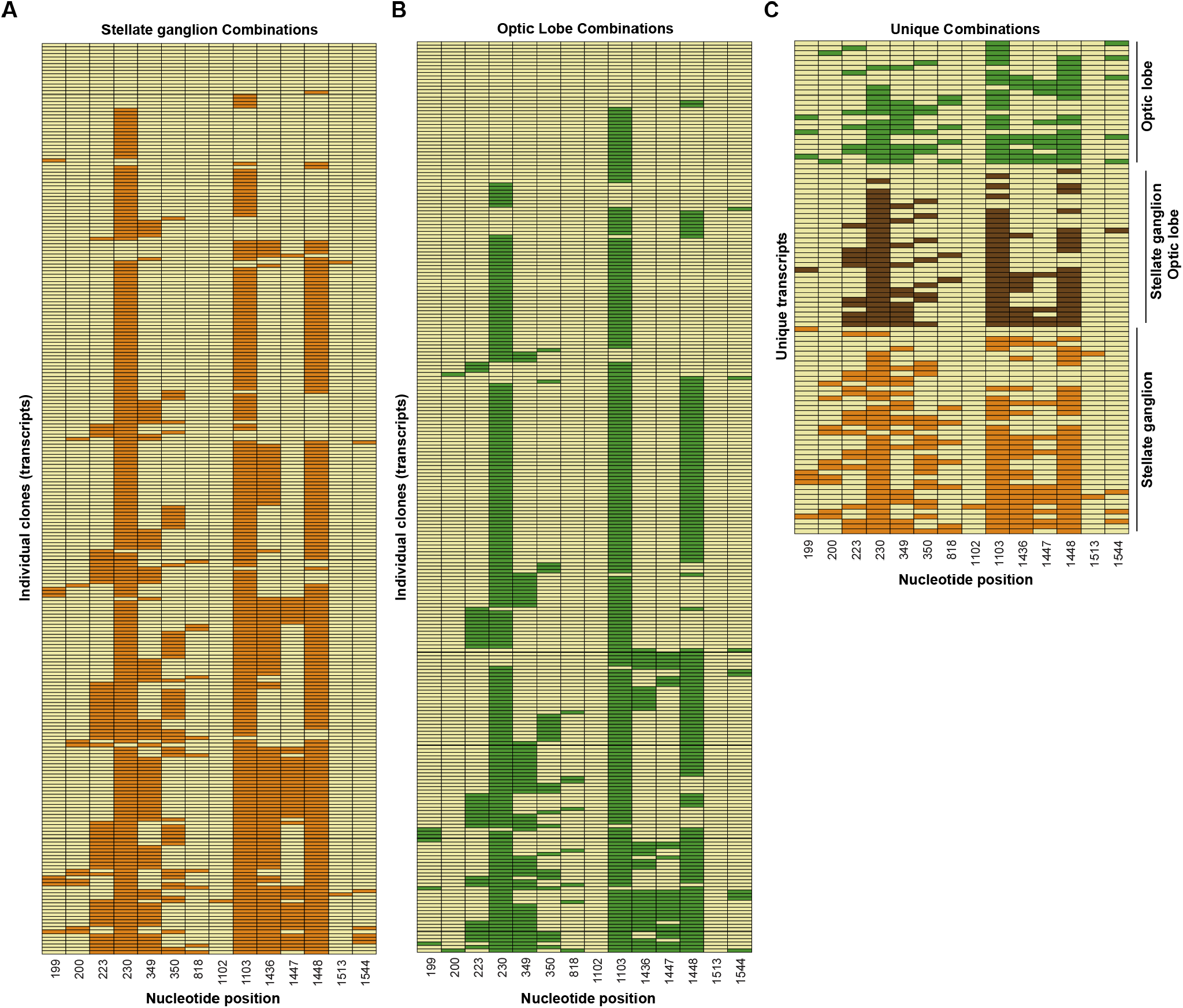
Summary of tissue-specific recoding in *D. pealeii*. Sequences of individual transcripts were determined by clone sequencing of cDNA extracted from **(A)** the stellate ganglion and **(B)** the optic lobe of *D. pealeii*. **(C)** Unique sequences detected from the stellate ganglion (orange) and the optic lobe (green). Sequences found in both tissues are shown in brown.

**Fig. S4.**
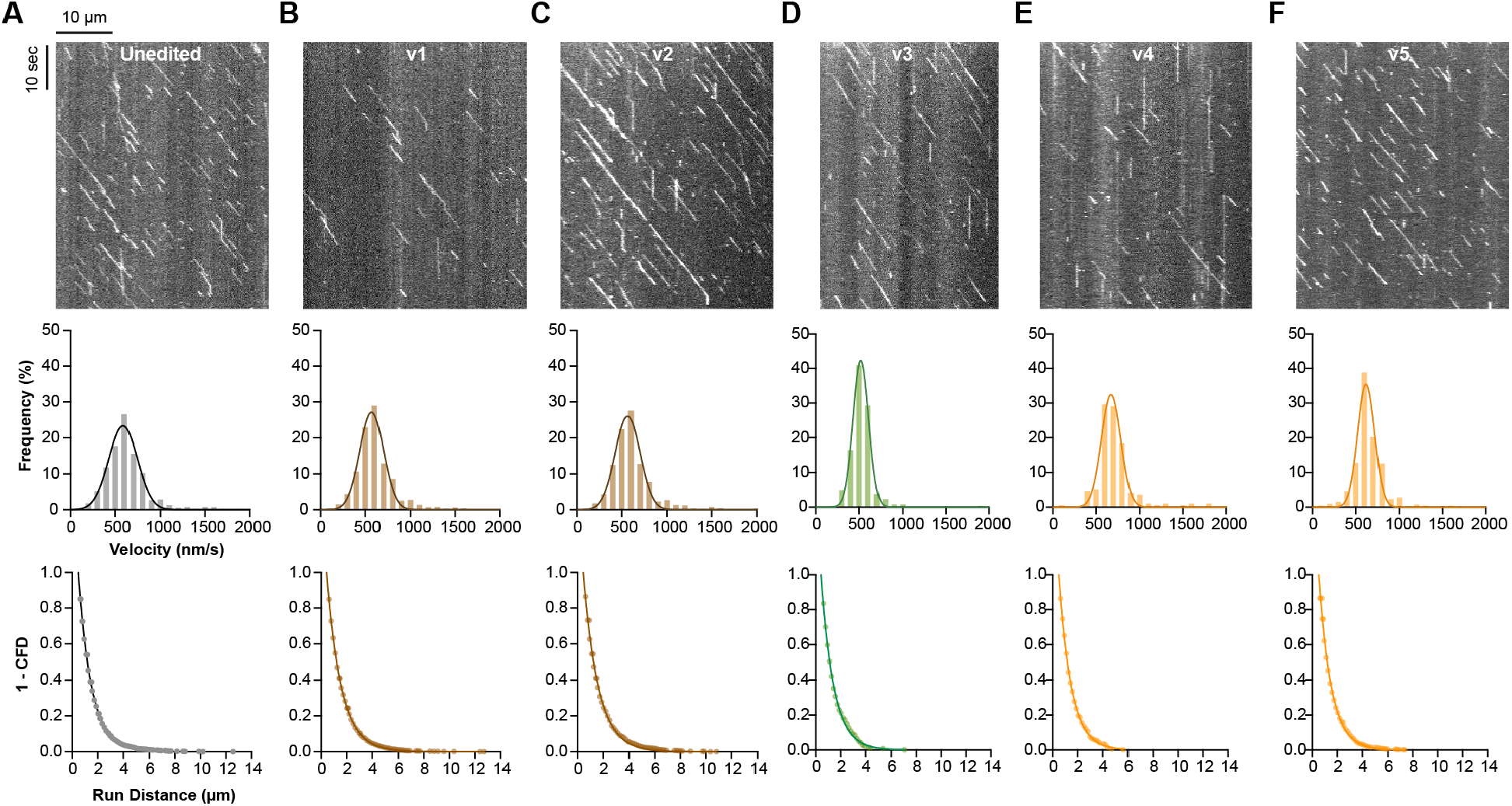
Summary of motility for *D. pealeii* kinesin variants. Representative kymographs, velocity frequency distributions and run distance frequency distributions for each *D. pealeii* kinesin (*Dpeal*K554) variant shown in Figure 2. Velocity histograms were fit to a Gaussian. For run distance measurements, 1-cumulative frequency distributions were fit to a one-phase exponential decay. See data S2 for velocity and run distance fit parameters including R^2^ and n for each condition. **(A)** Unedited **(B)** v1 **(C)** v2 **(D)** v3 **(E)** v4 **(F)** v5

**Fig. S5.**
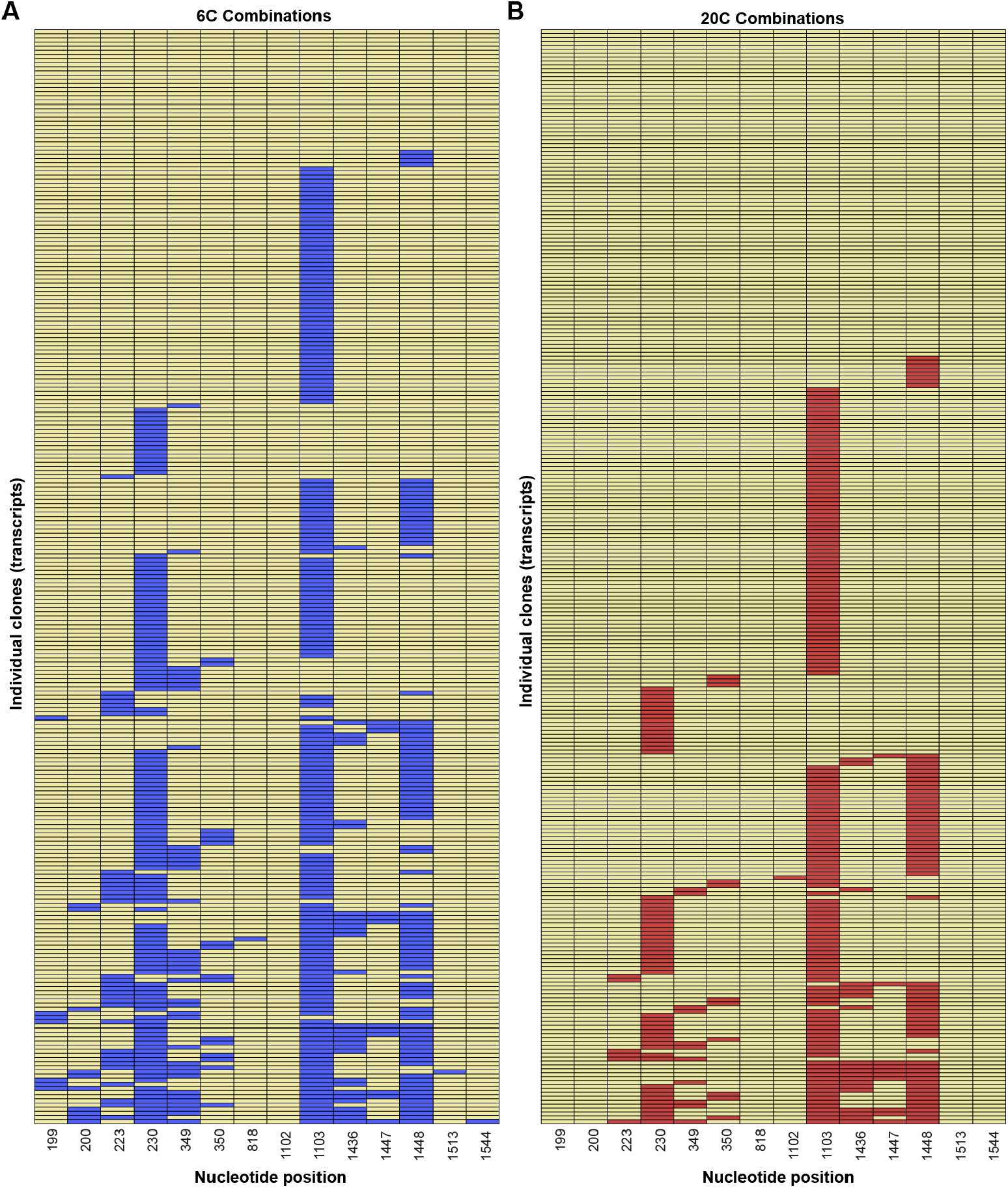
Summary of temperature-specific recoding in *D*.*opalescens*. Sequences of individual transcripts were determined by clone sequencing of cDNA extracted from *D. opalescens* exposed to **(A)** 6 °C water and **(B)** 20 °C water.

**Fig. S6.**
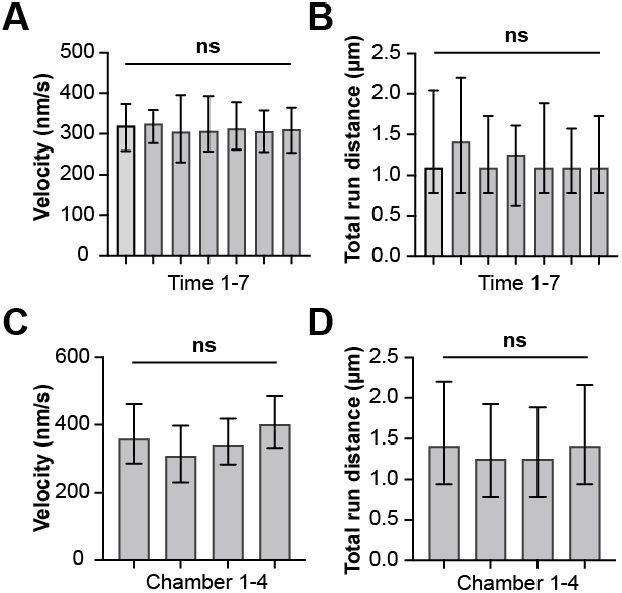
Controls for imaging of kinesin at 8 °C. **(A)** Median velocity +/- interquartile range and **(B)** Median run distance +/- interquartile range of human K560 at 8 °C in 1-minute videos taken sequentially. **(C)** Median velocity +/- interquartile range and b Median run distance +/- interquartile range of human K560 at 8 °C across different motility chambers.

**Fig. S7.**
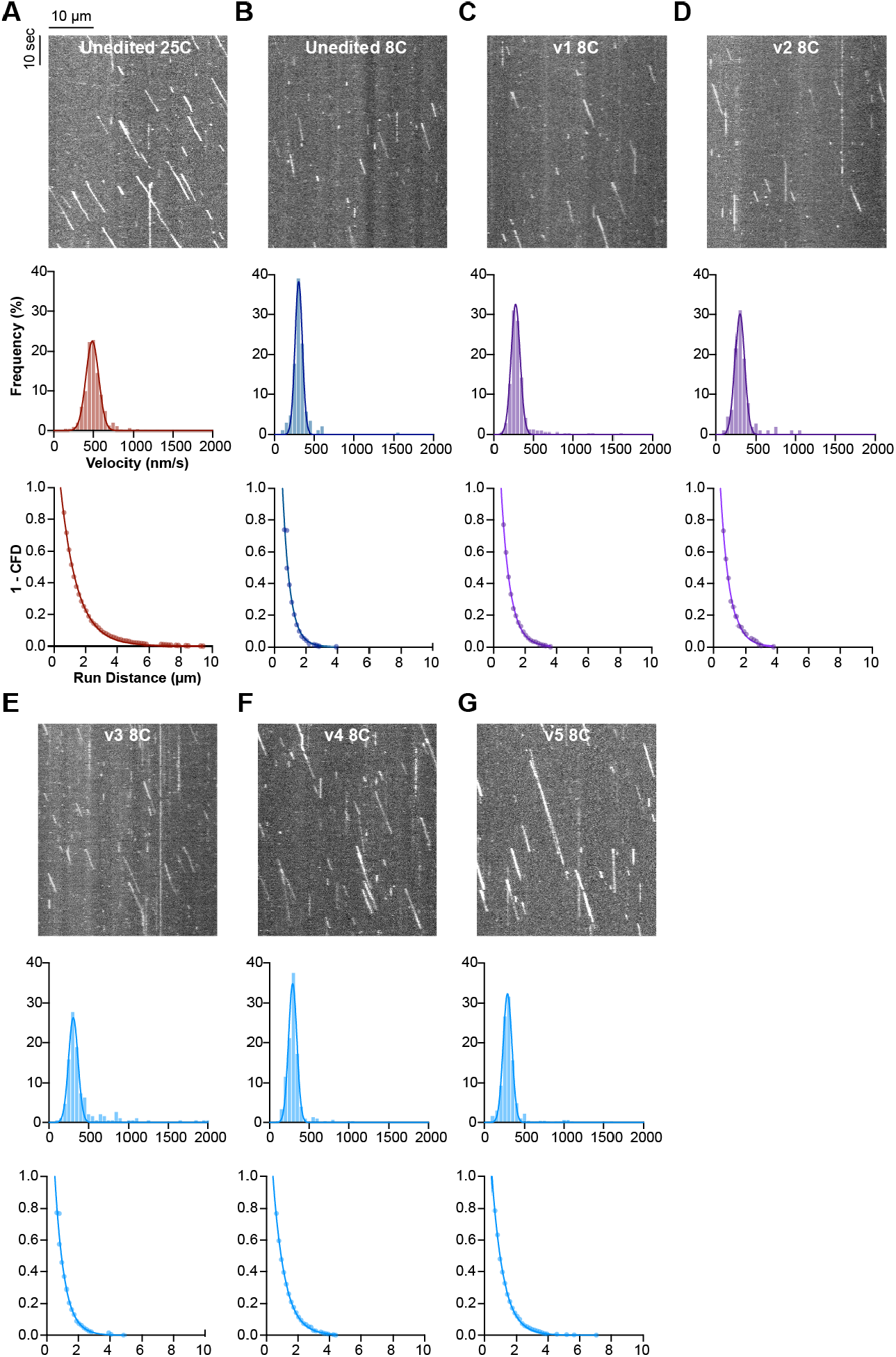
Summary of motility for *D. opalescens* kinesin variants at 8 °C. Representative kymographs, velocity frequency distributions and run distance frequency distributions for each *D. opalescens* kinesin (*Dopal*K554) variant shown in Figure 4. Velocity histograms were fit to a Gaussian. For run distance measurements, 1-cumulative frequency distributions were fit to a one-phase exponential decay. See data S2 for velocity and run distance fit parameters including R^2^ and n for each condition. **(A)** Unedited *D. opalescens* kinesin at 25 °C **(B)** Unedited *D. opalescens* kinesin at 8 °C **(C)** v1 **(D)** v2 **(E)** v3 **(F)** v4 **(G)** v5.

**Fig. S8.**
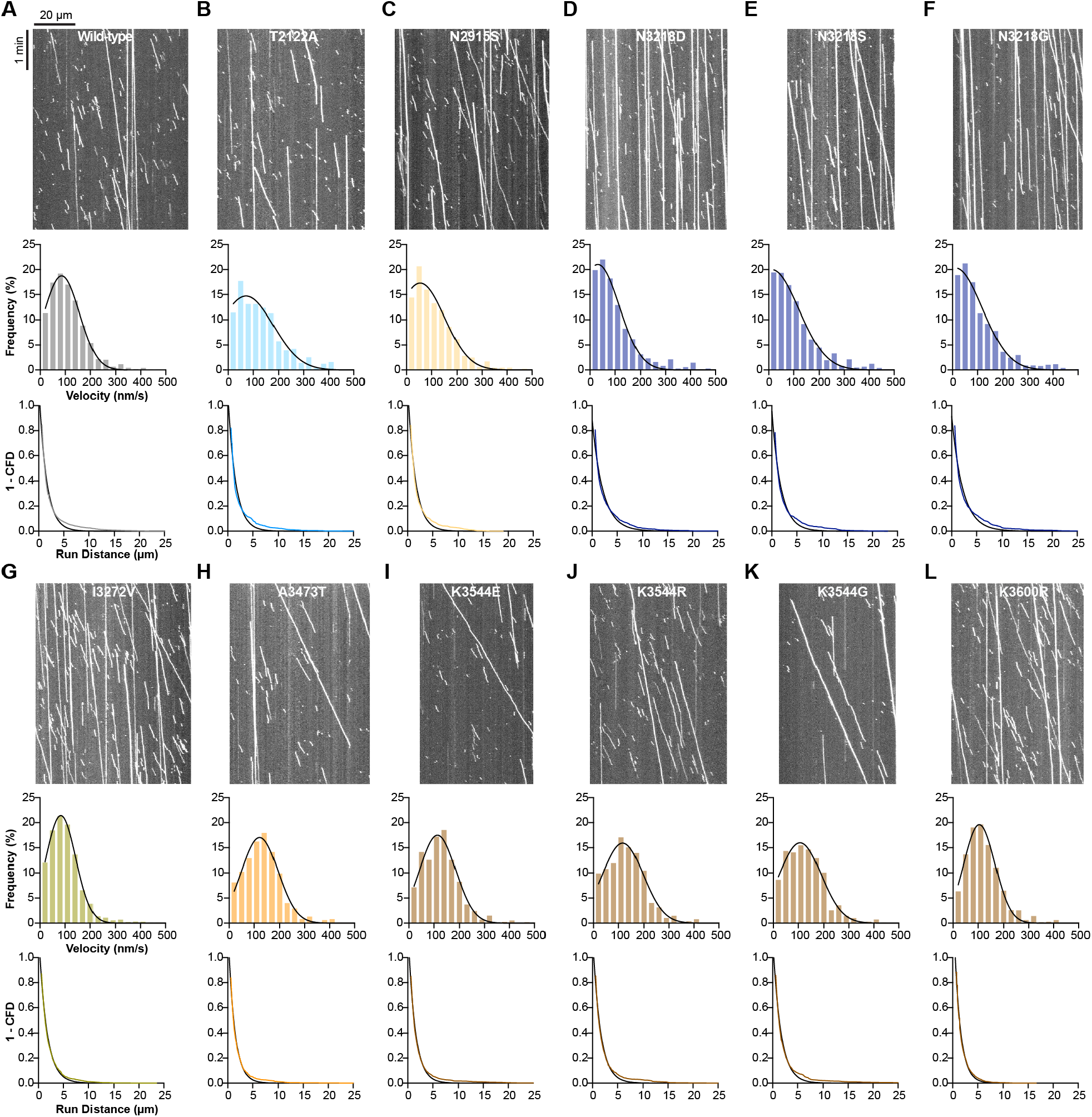
Summary of motility for yeast dynein mutants. Representative kymographs, velocity frequency distributions and run distance frequency distributions for each yeast dynein GST-Dyn1(331kDa) mutant in Figure 6. Velocity histograms were fit to a Gaussian. For run distance measurements, 1-cumulative frequency distributions were fit to a one-phase exponential decay. See data S2 for velocity and run distance fit parameters including R^2^ and n for each condition. **(A)** wild-type **(B)** T2122A **(C)** N2915S **(D)** N3218D **(E)** N3218S **(F)** N3218G **(G)** I3272V **(H)** A3473T **(I)** K3544E **(J)** K3544R **(K)** K3544G **(L)** K3600R.

**Fig. S9.**
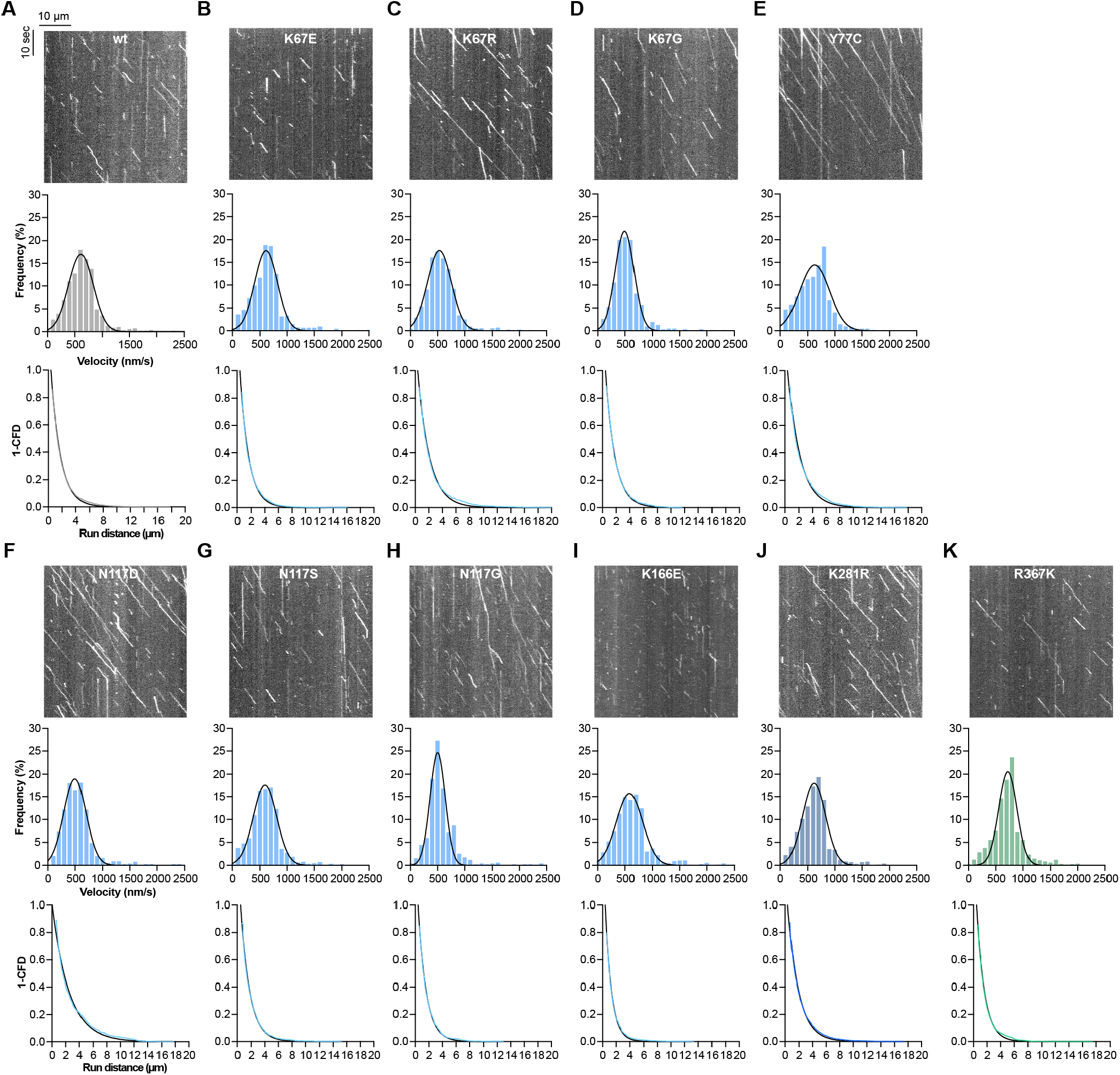
Summary of motility for human kinesin mutants. Representative kymographs, velocity frequency distributions and run distance frequency distributions for each human kinesin K560 mutant in Figure 6. Velocity histograms were fit to a Gaussian. For run distance measurements, 1-cumulative frequency distributions were fit to a one-phase exponential decay. See data S2 for velocity and run distance fit parameters including R^2^ and n for each condition. **(A)** wild-type **(B)** K67E **(C)** K67R **(D)** K67G **(E)** Y77C **(F)** N117D **(G)** N117S **(H)** N117G **(I)** K166E **(J)** K281R **(K)** R367K.

**Fig. S10.**
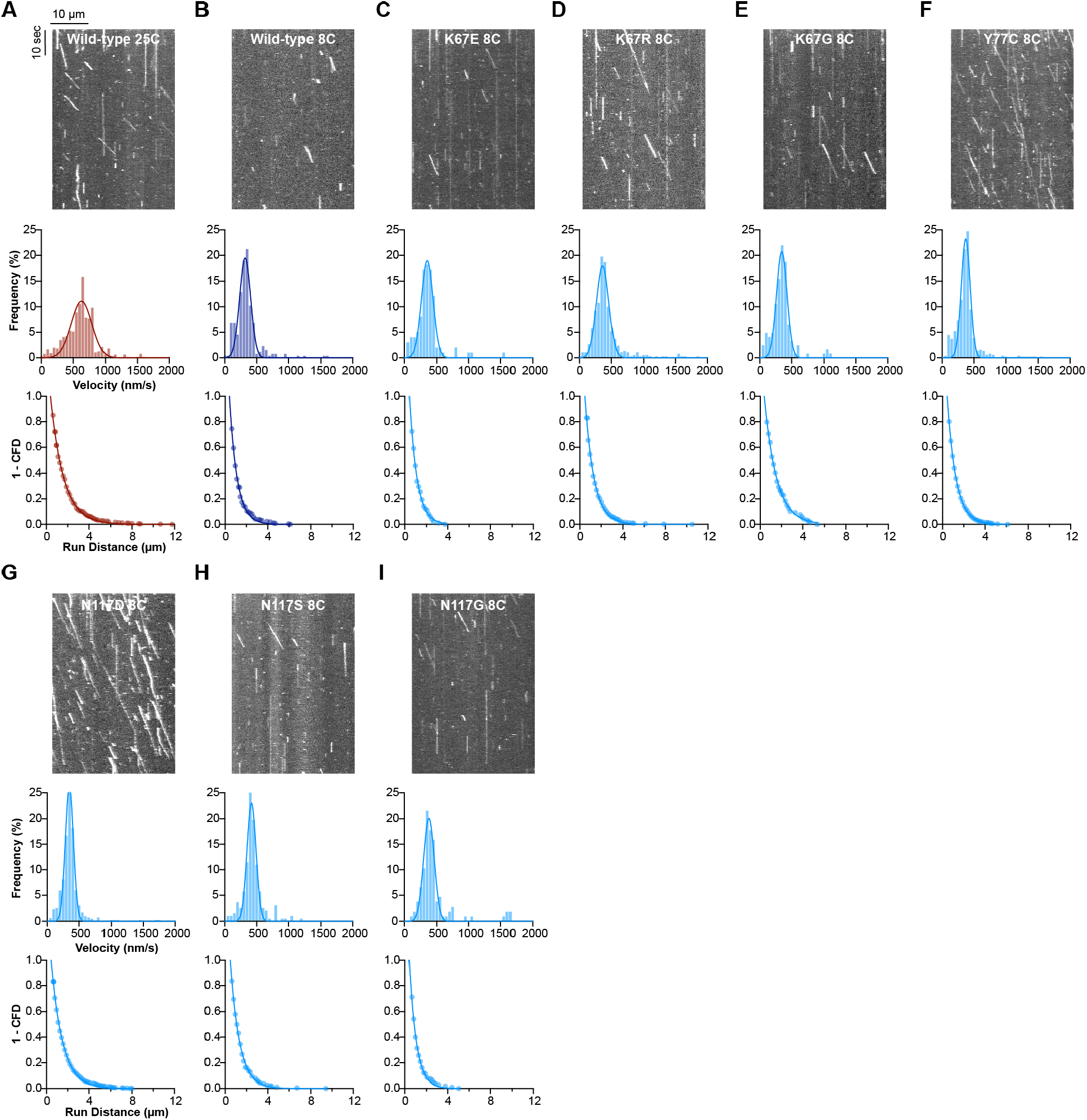
Summary of motility for human kinesin mutants at 8 °C. Representative kymographs, velocity frequency distributions and run distance frequency distributions for each human kinesin (K560-GFP) mutant shown in Figure 6. Velocity histograms were fit to a Gaussian. For run distance measurements, 1-cumulative frequency distributions were fit to a one-phase exponential decay. See data S2 for velocity and run distance fit parameters including R^2^ and n for each condition. **(A)** Wild-type at 25 °C **(B)** Wild-type at 8 °C **(C)** K67E **(D)** K67R **(E)** K67G **(F)** Y77C **(G)** N117D **(H)** N117S **(I)** N117G.

**Table S1.** Cephalopod recoding sites in cytoplasmic dynein. Number of combined cephalopod recoding sites detected across proteins in the cytoplasmic dynein transport complex (*4*).

| Name | Human homolog<br>(Uniprot Name) | *D. pealeii* | *S. officianalis* | *O. bimaculoides* | *O. vulgaris* | Sum |
| --- | --- | --- | --- | --- | --- | --- |
| **Dynein complex** |  |  |  |  |  |  |
| Cytoplasmic dynein 1 heavy chain 1 | DYHC1 | 87 | 194 | 78 | 129 | 488 |
| Cytoplasmic dynein 1 intermediate chain 2 | DC1I2 | 16 | 26 | 7 | 7 | 56 |
| Cytoplasmic dynein 1 light intermediate chain 2 | DC1L2 | 21 | 19 | 12 | 7 | 59 |
| Dynein light chain Tctex-type 1 | DYLT1 | 2 | 3 | 0 | 0 | 5 |
| No recoding sites found in: DYNLRB1, DYNLRB2, DYNLL1, DYNLL2, DC1I1, DC1L1, DYLT3 |  |  |  |  |  | Total: 608 |
| **Dynactin complex** |  |  |  |  |  |  |
| dynactin subunit 1/p150 | DCTN1 | 45 | 9 | 0 | 14 | 68 |
| dynactin subunit 2/p50 | DCTN2 | 9 | 8 | 6 | 6 | 29 |
| dynactin subunit 3/p22 | DCTN3 | 1 | 3 | 0 | 0 | 4 |
| ACTR1A/Arp1 | ACTZ | 8 | 2 | 2 | 5 | 17 |
| B-actin | ACTB | 6 | 2 | 9 | 29 | 46 |
| CapZA2 | CAPZA2 | 1 | 3 | 2 | 1 | 7 |
| CapZB | CAPZB | 3 | 0 | 0 | 1 | 4 |
| ACTR10/Arp11 | ARP10 | 8 | 11 | 11 | 14 | 44 |
| dynactin subunit 4/p62 | DCTN4 | 1 | 2 | 4 | 10 | 17 |
| dynactin subunit 4/p25 | DCTN5 | 0 | 1 | 0 | 1 | 2 |
| dynactin subunit 6/p27 | DCTN6 | 0 | 21 | 13 | 2 | 36 |
| No recoding sites found in: ACTY(ACTR1B), CAPZA1, CAPZA2 |  |  |  |  |  | Total: 274 |
| **Activating adaptors** |  |  |  |  |  |  |
| Protein bicaudal D homolog 1 | BICD1 | 20 | 23 | 30 | 35 | 108 |
| Trafficking kinesin-binding protein 1 | TRAK1 | 11 | 5 | 9 | 24 | 49 |
| BICD Family like cargo adaptor 1 | BICL1 | 0 | 4 | 1 | 6 | 11 |
| Hook homolog 3 | HOOK3 | 0 | 4 | 4 | 7 | 15 |
| Spindly | SPDLY | 0 | 10 | 3 | 0 | 13 |
| Rab 45 | RASEF | 1 | 0 | 0 | 0 | 1 |
| C-Jun-amino-terminal kinase-interacting protein 4 | JIP4 | 107 | 84 | 59 | 89 | 339 |
| No recoding sites found in: NINL, NIN, BICD2, BICDL2, HOOK1, HOOK2, RFIP3, CRACR2A, CCDC88A, CCDC88B, CCDC88C, JIP3, NUMA, TRAK2, HAP1 |  |  |  |  |  |  |
| **Other regulators:** |  |  |  |  |  |  |
| Platelet-activating factor acetylhydrolase IB subunit alpha | LIS1 | 4 | 5 | 15 | 25 | 49 |
| Nuclear distribution protein nudE homolog 1 | NDE1 | 26 | 10 | 2 | 1 | 39 |
| No recoding sites found in: NDEL1 |  |  |  |  |  |  |

**Table S2.** Cephalopod recoding sites in kinesins. Number of combined cephalopod recoding sites across kinesin family proteins (*4*).

| Kinesin family | Human homolog | *D. pealeii* | *S. officinalis* | *O. bimaculoides* | *O. vulgaris* | Sum |
| --- | --- | --- | --- | --- | --- | --- |
| kinesin-1 | KINH | 38 | 64 | 17 | 24 | 143 |
|  | KLC1 | 5 | 25 | 30 | 32 | 92 |
| kinesin-2 | KIF3A | 16 | 18 | 13 | 26 | 73 |
|  | KIF3B | 8 | 16 | 14 | 20 | 58 |
|  | KIFA3 | 83 | 64 | 11 | 41 | 199 |
| kinesin-3 | KIF1A | 63 | 72 | 0 | 45 | 180 |
|  | KIF1B | 0 | 0 | 25 | 0 | 25 |
|  | KI13A | 19 | 37 | 46 | 54 | 156 |
|  | KI13B | 5 | 8 | 0 | 0 | 13 |
|  | KIF14 | 0 | 4 | 7 | 18 | 29 |
|  | KI16B | 18 | 20 | 17 | 25 | 80 |
|  | KIF28 | 0 | 5 | 2 | 0 | 7 |
| kinesin-4 | KIF4A | 0 | 2 | 0 | 1 | 3 |
|  | KIF4B | 0 | 0 | 0 | 1 | 1 |
|  | KI21A | 12 | 5 | 12 | 10 | 39 |
|  | KI21B | 1 | 1 | 0 | 11 | 13 |
|  | KIF27 | 9 | 8 | 4 | 2 | 23 |
| kinesin-5 | KIF11 | 0 | 0 | 1 | 5 | 6 |
| kinesin-6 | KI20A | 0 | 10 | 3 | 2 | 15 |
|  | KIF23 | 0 | 0 | 0 | 2 | 2 |
| kinesin-8 | KI18A | 1 | 12 | 0 | 0 | 13 |
|  | KI18B | 1 | 0 | 0 | 0 | 1 |
| kinesin-9 | KIF6 | 0 | 7 | 0 | 1 | 8 |
|  | KIF9 | 0 | 0 | 5 | 6 | 11 |
| kinesin-10 | KIF22 | 0 | 4 | 0 | 0 | 4 |
| kinesin-11 | KI26B | 0 | 0 | 4 | 22 | 26 |
| kinesin-12 | KIF12 | 0 | 0 | 1 | 0 | 1 |
| kinesin-13 | KIF2A | 9 | 13 | 1 | 4 | 27 |
|  | KIF24 | 22 | 21 | 7 | 6 | 56 |
| kinesin-14a | KIFC1 | 0 | 0 | 0 | 3 | 3 |
| kinesin-14b | KIFC3 | 8 | 21 | 0 | 10 | 39 |
|  | KIF25 | 0 | 0 | 0 | 18 | 18 |

**Data S1. (separate file)**

Top 100 GO terms (category: biological process) enriched in *D. pealeii* edited transcripts (highlighted in blue) or unedited transcripts (highlighted in yellow) as compared to the full *D. pealeii* transcriptome.

**Data S2. (separate file)**

Summary of velocity and run distance fit parameters for single-molecule motility assays.

**Data S3. (separate file)**

Cephalopod RNA recoding sites in DYNC1H1.

**Data S4. (separate file)**

Cephalopod RNA recoding sites in KIF5B.

## References

1. K. Nishikura, A-to-I editing of coding and non-coding RNAs by ADARs. Nat. Rev. Mol. Cell Biol. 17, 83–96 (2016).

2. O. Gabay, Y. Shoshan, E. Kopel, U. Ben-Zvi, T. D. Mann, N. Bressler, R. Cohen-Fultheim, A. A. Schaffer, S. H. Roth, Z. Tzur, E. Y. Levanon, E. Eisenberg, Landscape of adenosine-to-inosine RNA recoding across human tissues. Nat. Commun. 13, 1184 (2022).

3. E. Eisenberg, Proteome Diversification by RNA Editing. Methods Mol. Biol. 2181, 229–251 (2021).

4. N. Liscovitch-Brauer, S. Alon, H. T. Porath, B. Elstein, R. Unger, T. Ziv, A. Admon, E. Y. Levanon, J. J. C. Rosenthal, E. Eisenberg, Trade-off between Transcriptome Plasticity and Genome Evolution in Cephalopods. Cell. 169, 191-202.e11 (2017).

5. S. Alon, S. C. Garrett, E. Y. Levanon, S. Olson, B. R. Graveley, J. J. C. Rosenthal, E. Eisenberg, The majority of transcripts in the squid nervous system are extensively recoded by A-to-I RNA editing. eLife. 4, e05198 (2015).

6. I. C. Vallecillo-Viejo, N. Liscovitch-Brauer, J. F. Diaz Quiroz, M. F. Montiel-Gonzalez, S. E. Nemes, K. J. Rangan, S. R. Levinson, E. Eisenberg, J. J. C. Rosenthal, Spatially regulated editing of genetic information within a neuron. Nucleic Acids Res. 48, 3999–4012 (2020).

7. S. Garrett, J. J. C. Rosenthal, RNA editing underlies temperature adaptation in K+ channels from polar octopuses. Science. 335, 848–851 (2012).

8. J. J. C. Rosenthal, The emerging role of RNA editing in plasticity. J. Exp. Biol. 218, 1812–1821 (2015).

9. C. B. Albertin, S. Medina-Ruiz, T. Mitros, H. Schmidbaur, G. Sanchez, Z. Y. Wang, J. Grimwood, J. J. C. Rosenthal, C. W. Ragsdale, O. Simakov, D. S. Rokhsar, Genome and transcriptome mechanisms driving cephalopod evolution. Nat. Commun. 13, 2427 (2022).

10. M. Moldovan, Z. Chervontseva, G. Bazykin, M. S. Gelfand, Adaptive evolution at mRNA editing sites in soft-bodied cephalopods. PeerJ. 8, e10456 (2020).

11. Y. Shoshan, N. Liscovitch-Brauer, J. J. C. Rosenthal, E. Eisenberg, Adaptive Proteome Diversification by Nonsynonymous A-to-I RNA Editing in Coleoid Cephalopods. Mol. Biol. Evol. 38, 3775–3788 (2021).

12. The preponderance of nonsynonymous A-to-I RNA editing in coleoids is nonadaptive. Nat. Commun. 10, 5411 (2019).

13. C. Colina, J. P. Palavicini, D. Srikumar, M. Holmgren, J. J. C. Rosenthal, Regulation of Na+/K+ ATPase transport velocity by RNA editing. PLoS Biol. 8, e1000540 (2010).

14. J. J. C. Rosenthal, F. Bezanilla, Extensive editing of mRNAs for the squid delayed rectifier K+ channel regulates subunit tetramerization. Neuron. 34, 743–757 (2002).

15. D. E. Patton, T. Silva, F. Bezanilla, RNA editing generates a diverse array of transcripts encoding squid Kv2 K+ channels with altered functional properties. Neuron. 19, 711–722 (1997).

16. M. Holmgren, J. J. C. Rosenthal, Regulation of Ion Channel and Transporter Function Through RNA Editing. Curr. Issues Mol. Biol. 17, 23–36 (2015).

17. L. Zeidberg, Advances in Squid Biology, Ecology and Fisheries. Part I – Myopsid Squids. pp. 159–204 (2013).

18. L. D. Jacobson, Essential fish habitat source document. Longfin inshore squid, Loligo pealeii, life history and habitat characteristics. NOAA Tech. Memo. NMFS-NE. 193 (2005).

19. K. Kawaguchi, S. Ishiwata, Temperature dependence of force, velocity, and processivity of single kinesin molecules. Biochem. Biophys. Res. Commun. 272, 895–899 (2000).

20. F. Doval, K. Chiba, R. J. McKenney, K. M. Ori-McKenney, M. D. Vershinin, Temperature-dependent activity of kinesins is regulable. Biochem. Biophys. Res. Commun. 528, 528–530 (2020).

21. W. Hong, A. Takshak, O. Osunbayo, A. Kunwar, M. Vershinin, The Effect of Temperature on Microtubule-Based Transport by Cytoplasmic Dynein and Kinesin-1 Motors. Biophys. J. 111, 1287–1294 (2016).

22. L. D. Zeidberg, G. Isaac, C. L. Widmer, H. Neumeister, W. F. Gilly, Egg capsule hatch rate and incubation duration of the California market squid, Doryteuthis (= Loligo) opalescens: insights from laboratory manipulations. Mar. Ecol. 32, 468–479 (2011).

23. A. P. Carter, A. G. Diamant, L. Urnavicius, How dynein and dynactin transport cargos: a structural perspective. Curr. Opin. Struct. Biol. 37, 62–70 (2016).

24. G. Bhabha, G. T. Johnson, C. M. Schroeder, R. D. Vale, How Dynein Moves Along Microtubules. Trends Biochem. Sci. 41, 94–105 (2016).

25. W. O. Hancock, The Kinesin-1 Chemomechanical Cycle: Stepping Toward a Consensus. Biophys. J. 110, 1216–1225 (2016).

26. T. Kon, T. Oyama, R. Shimo-Kon, K. Imamula, T. Shima, K. Sutoh, G. Kurisu, The 2.8 Å crystal structure of the dynein motor domain. Nature. 484, 345–350 (2012).

27. H. Schmidt, E. S. Gleave, A. P. Carter, Insights into dynein motor domain function from a 3.3-Å crystal structure. Nat. Struct. Mol. Biol. 19, 492–497, S1 (2012).

28. A. J. Roberts, B. Malkova, M. L. Walker, H. Sakakibara, N. Numata, T. Kon, R. Ohkura, T. A. Edwards, P. J. Knight, K. Sutoh, K. Oiwa, S. A. Burgess, ATP-Driven Remodeling of the Linker Domain in the Dynein Motor. Structure. 20, 1670–1680 (2012).

29. K. Zhang, H. E. Foster, A. Rondelet, S. E. Lacey, N. Bahi-Buisson, A. W. Bird, A. P. Carter, Cryo-EM Reveals How Human Cytoplasmic Dynein Is Auto-inhibited and Activated. Cell. 169, 1303-1314.e18 (2017).

30. F. J. Kull, E. P. Sablin, R. Lau, R. J. Fletterick, R. D. Vale, Crystal structure of the kinesin motor domain reveals a structural similarity to myosin. Nature. 380, 550–555 (1996).

31. S. Rice, A. W. Lin, D. Safer, C. L. Hart, N. Naber, B. O. Carragher, S. M. Cain, E. Pechatnikova, E. M. Wilson-Kubalek, M. Whittaker, E. Pate, R. Cooke, E. W. Taylor, R. A. Milligan, R. D. Vale, A structural change in the kinesin motor protein that drives motility. Nature. 402, 778–784 (1999).

32. H. Y. K. Kaan, D. D. Hackney, F. Kozielski, The structure of the kinesin-1 motor-tail complex reveals the mechanism of autoinhibition. Science. 333, 883–885 (2011).

33. K. A. Dietrich, C. V. Sindelar, P. D. Brewer, K. H. Downing, C. R. Cremo, S. E. Rice, The kinesin-1 motor protein is regulated by a direct interaction of its head and tail. Proc. Natl. Acad. Sci. 105, 8938–8943 (2008).

34. W. Hwang, M. J. Lang, M. Karplus, Force Generation in Kinesin Hinges on Cover-Neck Bundle Formation. Structure. 16, 62–71 (2008).

35. B. G. Budaitis, S. Jariwala, L. Rao, Y. Yue, D. Sept, K. J. Verhey, A. Gennerich, Pathogenic mutations in the kinesin-3 motor KIF1A diminish force generation and movement through allosteric mechanisms. J. Cell Biol. 220, e202004227 (2021).

36. G. Woehlke, A. K. Ruby, C. L. Hart, B. Ly, N. Hom-Booher, R. D. Vale, Microtubule Interaction Site of the Kinesin Motor. Cell. 90, 207–216 (1997).

37. K. Poirier, N. Lebrun, L. Broix, G. Tian, Y. Saillour, C. Boscheron, E. Parrini, S. Valence, B. S. Pierre, M. Oger, D. Lacombe, D. Geneviève, E. Fontana, F. Darra, C. Cances, M. Barth, D. Bonneau, B. D. Bernadina, S. N’guyen, C. Gitiaux, P. Parent, V. des Portes, J. M. Pedespan, V. Legrez, L. Castelnau-Ptakine, P. Nitschke, T. Hieu, C. Masson, D. Zelenika, A. Andrieux, F. Francis, R. Guerrini, N. J. Cowan, N. Bahi-Buisson, J. Chelly, Mutations in TUBG1, DYNC1H1, KIF5C and KIF2A cause malformations of cortical development and microcephaly. Nat. Genet. 45, 639–647 (2013).

38. S. L. Reck-Peterson, A. Yildiz, A. P. Carter, A. Gennerich, N. Zhang, R. D. Vale, Single-Molecule Analysis of Dynein Processivity and Stepping Behavior. Cell. 126, 335–348 (2006).

39. R. D. Vale, T. Funatsu, D. W. Pierce, L. Romberg, Y. Harada, T. Yanagida, Direct observation of single kinesin molecules moving along microtubules. Nature. 380, 451–453 (1996).

40. H. T. Hoang, M. A. Schlager, A. P. Carter, S. L. Bullock, DYNC1H1 mutations associated with neurological diseases compromise processivity of dynein–dynactin–cargo adaptor complexes. Proc. Natl. Acad. Sci. 114, E1597–E1606 (2017).

41. M. G. Marzo, J. M. Griswold, K. M. Ruff, R. E. Buchmeier, C. P. Fees, S. M. Markus, Molecular basis for dyneinopathies reveals insight into dynein regulation and dysfunction. eLife. 8, e47246 (2019).

42. S. Jennings, M. Chenevert, L. Liu, M. Mottamal, E. J. Wojcik, T. M. Huckaba, Characterization of kinesin switch I mutations that cause hereditary spastic paraplegia. PLoS ONE. 12, e0180353 (2017).

43. L. Rao, F. Berger, M. P. Nicholas, A. Gennerich, Molecular mechanism of cytoplasmic dynein tension sensing. Nat. Commun. 10, 3332 (2019).

44. W. B. Redwine, R. Hernandez-Lopez, S. Zou, J. Huang, S. L. Reck-Peterson, A. E. Leschziner, Structural basis for microtubule binding and release by dynein. Science. 337, 1532–1536 (2012).

45. S. E. Lacey, S. He, S. H. Scheres, A. P. Carter, Cryo-EM of dynein microtubule-binding domains shows how an axonemal dynein distorts the microtubule. eLife. 8, e47145 (2019).

46. B. J. Grant, D. M. Gheorghe, W. Zheng, M. Alonso, G. Huber, M. Dlugosz, J. A. McCammon, R. A. Cross, Electrostatically Biased Binding of Kinesin to Microtubules. PLOS Biol. 9, e1001207 (2011).

47. A. L. Yablonovitch, P. Deng, D. Jacobson, J. B. Li, The evolution and adaptation of A-to-I RNA editing. PLoS Genet. 13, e1007064 (2017).

48. L. E. Rieder, Y. A. Savva, M. A. Reyna, Y.-J. Chang, J. S. Dorsky, A. Rezaei, R. A. Reenan, Dynamic response of RNA editing to temperature in Drosophila. BMC Biol. 13, 1 (2015).

49. I. Buchumenski, O. Bartok, R. Ashwal-Fluss, V. Pandey, H. T. Porath, E. Y. Levanon, S. Kadener, Dynamic hyper-editing underlies temperature adaptation in Drosophila. PLOS Genet. 13, e1006931 (2017).

50. K. A. Riemondy, A. E. Gillen, E. A. White, L. K. Bogren, J. R. Hesselberth, S. L. Martin, Dynamic temperature-sensitive A-to-I RNA editing in the brain of a heterothermic mammal during hibernation. RNA. 24, 1481–1495 (2018).

51. Y. Duan, S. Dou, S. Luo, H. Zhang, J. Lu, Adaptation of A-to-I RNA editing in Drosophila. PLOS Genet. 13, e1006648 (2017).

52. T. A. Nguyen, J. W. J. Heng, P. Kaewsapsak, E. P. L. Kok, D. Stanojevic, H. Liu, A. Cardilla, A. Praditya, Z. Yi, M. Lin, J. G. A. Aw, Y. Y. Ho, K. L. E. Peh, Y. Wang, Q. Zhong, J. Heraud-Farlow, S. Xue, B. Reversade, C. Walkley, Y. S. Ho, M. Šikic, Y. Wan, M. H. Tan, Direct identification of A-to-I editing sites with nanopore native RNA sequencing. Nat. Methods. 19, 833–844 (2022).

53. A. M. Wenger, P. Peluso, W. J. Rowell, P.-C. Chang, R. J. Hall, G. T. Concepcion, J. Ebler, A. Fungtammasan, A. Kolesnikov, N. D. Olson, A. Töpfer, M. Alonge, M. Mahmoud, Y. Qian, C.-S. Chin, A. M. Phillippy, M. C. Schatz, G. Myers, M. A. DePristo, J. Ruan, T. Marschall, F. J. Sedlazeck, J. M. Zook, H. Li, S. Koren, A. Carroll, D. R. Rank, M. W. Hunkapiller, Accurate circular consensus long-read sequencing improves variant detection and assembly of a human genome. Nat. Biotechnol. 37, 1155–1162 (2019).

54. C. Notredame, D. G. Higgins, J. Heringa, T-coffee: a novel method for fast and accurate multiple sequence alignment 1 1Edited by J. Thornton. J. Mol. Biol. 302, 205–217 (2000).

55. J. Huang, A. J. Roberts, A. E. Leschziner, S. L. Reck-Peterson, Lis1 Acts as a “Clutch” between the ATPase and Microtubule-Binding Domains of the Dynein Motor. Cell. 150, 975–986 (2012).

56. A. J. Roberts, B. S. Goodman, S. L. Reck-Peterson, Reconstitution of dynein transport to the microtubule plus end by kinesin. eLife. 3, e02641 (2014).

57. S. Berg, D. Kutra, T. Kroeger, C. N. Straehle, B. X. Kausler, C. Haubold, M. Schiegg, J. Ales, T. Beier, M. Rudy, K. Eren, J. I. Cervantes, B. Xu, F. Beuttenmueller, A. Wolny, C. Zhang, U. Koethe, F. A. Hamprecht, A. Kreshuk, ilastik: interactive machine learning for (bio)image analysis. Nat. Methods. 16, 1226–1232 (2019).

58. G. Bhabha, H.-C. Cheng, N. Zhang, A. Moeller, M. Liao, J. A. Speir, Y. Cheng, R. D. Vale, Allosteric Communication in the Dynein Motor Domain. Cell. 159, 857–868 (2014).

59. M. G. Marzo, J. M. Griswold, S. M. Markus, Pac1/LIS1 stabilizes an uninhibited conformation of dynein to coordinate its localization and activity. Nat. Cell Biol. 22, 559–569 (2020).

60. B. G. Budaitis, S. Jariwala, D. N. Reinemann, K. I. Schimert, G. Scarabelli, B. J. Grant, D. Sept, M. J. Lang, K. J. Verhey, Neck linker docking is critical for Kinesin-1 force generation in cells but at a cost to motor speed and processivity. eLife. 8, e44146 (2019).

61. M. Dutta, M. R. Diehl, J. N. Onuchic, B. Jana, Structural consequences of hereditary spastic paraplegia disease-related mutations in kinesin. Proc. Natl. Acad. Sci. 115, E10822–E10829 (2018).

